# How to control for confounds in decoding analyses of neuroimaging data

**DOI:** 10.1101/290684

**Authors:** Lukas Snoek, Steven Miletić, H. Steven Scholte

## Abstract

Over the past decade, multivariate pattern analyses and especially decoding analyses have become a popular alternative to traditional mass-univariate analyses in neuroimaging research. However, a fundamental limitation of decoding analyses is that the source of information driving the decoder is ambiguous, which becomes problematic when the to-be-decoded variable is confounded by variables that are not of primary interest. In this study, we use a comprehensive set of simulations and analyses of empirical data to evaluate two techniques that were previously proposed and used to control for confounding variables in decoding analyses: counterbalancing and confound regression. For our empirical analyses, we attempt to decode gender from structural MRI data when controlling for the confound “brain size”. We show that both methods introduce strong biases in decoding performance: counterbalancing leads to better performance than expected (i.e., positive bias), which we show in our simulations is due to the subsampling process that tends to remove samples that are hard to classify; confound regression, on the other hand, leads to worse performance than expected (i.e., negative bias), even resulting in significant below-chance performance in some scenarios. In our simulations, we show that below-chance accuracy can be predicted by the variance of the distribution of correlations between the features and the target. Importantly, we show that this negative bias disappears in both the empirical analyses and simulations when the confound regression procedure performed in every fold of the cross-validation routine, yielding plausible model performance. From these results, we conclude that foldwise confound regression is the only method that appropriately controls for confounds, which thus can be used to gain more insight into the exact source(s) of information driving one’s decoding analysis.

**HIGHLIGHTS:**

- The interpretation of decoding models is ambiguous when dealing with confounds;
- We evaluate two methods, counterbalancing and confound regression, in their ability to control for confounds;
- We find that counterbalancing leads to positive bias because it removes hard-to-classify samples;
- We find that confound regression leads to negative bias, because it yields data with less signal than expected by chance;
- Our simulations demonstrate a tight relationship between model performance in decoding analyses and the sample distribution of the correlation coefficient;
- We show that the negative bias observed in confound regression can be remedied by cross-validating the confound regression procedure.

## INTRODUCTION

In the past decade, multivariate pattern analyses (MVPA) have emerged as a popular alternative to traditional univariate analyses of neuroimaging data (Haxby, 2012; Norman, Polyn, Detre, & Haxby, 2006). The defining feature of MVPA is that it considers *patterns* of brain activation instead of single units of activation (i.e., voxels in MRI, sensors in MEG/EEG). One of the most-often used MVPA methods is “decoding”, in which machine learning algorithms are applied to neuroimaging data directly to decode a particular stimulus, task, or psychometric feature. For example, decoding analyses have been used to successfully decode different experimental conditions within subjects, such as object category from fMRI activity patterns (Haxby et al., 2001) and working memory representations from EEG data (LaRocque, Lewis-Peacock, Drysdale, Oberauer, & Postle, 2013), as well between-subject factors such as Alzheimer’s disease (vs. healthy controls) from structural MRI data (Cuingnet et al., 2011) and major depressive disorder (vs. healthy controls) from resting-state functional connectivity (Craddock, Holtzheimer, Hu, & Mayberg, 2009). One reason for the popularity of MVPA methods, and especially decoding, is that they appear to be more sensitive than traditional mass-univariate methods in detecting effects of interest, which is often attributed to the ability of MVPA to pick up spatially distributed multidimensional representations while univariate methods, by definition, cannot (Jimura & Poldrack, 2012).

In the past years, however, MVPA’s apparent superior sensitivity and ability to pick up distributed representations has been criticized for a number of reasons, both statistical (Allefeld, Görgen, & Haynes, 2016; Davis et al., 2014; Gilron, Rosenblatt, Koyejo, Poldrack, & Mukamel, 2017; Haufe et al., 2014) and more conceptual (Naselaris & Kay, 2015; Weichwald et al., 2015) in nature. For the purposes of the current study, we focus on the specific criticism forwarded by Naselaris and Kay (2015), who argue that decoding analyses are inherently “representationally ambiguous” (see Popov, Ostarek, & Tenison, 2018, for a similar argument in the context of encoding analyses). This representational ambiguity arises when the classes of the to-be-decoded variable systematically vary in more than one source of information (see also Carlson & Wardle, 2015; Ritchie, Kaplan, & Klein, 2017; Weichwald et al., 2015). The current study aims to investigate how decoding analyses can be made more interpretable by reducing representational ambiguity.

To illustrate the problem of representational ambiguity, consider, for example, the scenario in which a researcher aims to decode gender (male/female) from structural MRI with the aim to contribute to the understanding of gender differences — an endeavour that generated considerable interest and controversy within the scientific community (Chekroud, Ward, Rosenberg, & Holmes, 2016; Del Giudice et al., 2016; Glezerman, 2016; Joel & Fausto-Sterling, 2016; Rosenblatt, 2016). By performing a decoding analysis on the MRI data, the researcher hopes to capture meaningful patterns of variation in the data of male and female participants that are predictive of the participant’s gender. From the literature, we know that gender dimorphism in the brain is manifested in two major ways (Good et al., 2001; O’Brien et al., 2011). First, there is a *global* difference between male and female brains: men have on average about 15% larger intracranial volume than women, which falls in the range of gender differences in height (8.2%) and weight (18.7%; Gur et al., 1999; Lüders, Steinmetz, & Jäncke, 2002). Second, brains of men and women are known to differ *locally*: some specific brain areas are found to be larger in women than in men (e.g., in superior and middle temporal cortex; Good et al., 2001) and vice versa (e.g., in frontomedial cortex; Goldstein et al., 2001). One could argue that, given that one is interested in explaining behavioral or mental gender differences, global (i.e., absolute) differences are relatively uninformative, as it reflects the fact than male *bodies* are in general larger than female bodies (Gur et al., 1999; Sepehrband et al., 2018). As such, our hypothetical researcher is likely primarily interested in the *local* sources of variation in the neuroanatomy of male and female brains.

Now, suppose that the researcher is able to decode gender from the MRI data significantly above chance, it remains unclear on which source of information the decoder is capitalizing: the (arguably meaningful) local difference in brain structure or the (in the context of this question arguably uninteresting) global difference in brain size? In other words, the data contains more than one source of information that may be used to predict gender. In the current study, we aim to evaluate methods that improve the interpretability of decoding analyses through controlling for these “uninteresting” sources of information.

### Partitioning effects into *true* signal and *confounded* signal

Are multiple sources of information necessarily problematic? And what makes a source of information interesting or uninteresting? First of all, this depends on the particular goal of the researcher employing the decoding analysis. In principle, multiple sources of information in the data does not pose a problem if a researcher is only interested in accurate *prediction* and not necessarily *interpretability* of the model (Bzdok, 2017; Haufe et al., 2014; Hebart & Baker, 2017). In brain-computer interfaces (BCI), for example, accurate *prediction* is arguably relatively more important than *interpretability*, i.e., knowing which sources of information are driving the decoder. Similarly, if the researcher from our gender-decoding example is only interested in accurately predicting gender regardless of model interpretability, representational ambiguity is not a problem^1^. As such, whether representational ambiguity in decoding analyses is a problem thus depends on the specific goal of the researcher (Hebart & Baker, 2017).

In most scientific applications of decoding analyses, however, model interpretability is important, because researchers are often interested in the relative contributions of different sources of information in their models. Specifically, in most decoding analyses, researchers often (implicitly) assume that the decoder is *only* using information in the neuroimaging data that is related to the variable that is being decoded (Ritchie, Kaplan, & Klein, 2017). In this scenario, representational ambiguity (i.e., the presence of *multiple* sources of information) *is* problematic as it violates the (implicit) assumption that the decoded variable is the *only* source of information driving the decoder. Another way conceptualize the problem of representational ambiguity is that, using the aforementioned example, (global) brain size is *confounding* the decoding analysis of gender. Here, we define a confound as *a variable that is not of primary interest, which correlates with the to-be-decoded variable (the target) and is encoded in the imaging data^2^*.

To give an example of confounding variables in the context of decoding, suppose one is interested in building a classifier that is able to predict whether subjects are suffering from schizophrenia or not based on the subjects’ gray matter data. Here, the variable “schizophrenia-or-not” is the variable of interest, which is assumed to be encoded in the neuroimaging data (i.e., the gray matter) and can thus be decoded. However, there are multiple factors known to covary with schizophrenia, like gender (i.e., men are more often afflicted than women; McGrath, Saha, Chant, & Welham, 2008) and substance abuse (Dixon, 1999), which are also known to affect gray matter (Bangalore et al., 2008; Gur et al., 1999; Van Haren, Cahn, Hulshoff Pol, & Kahn, 2013). As such, the variables gender and substance abuse can be considered confounds according to our definition, because they are both correlated with the target (schizophrenia or not) and are known to be encoded in the neuroimaging data (i.e., the effect of these variables is represented in the gray matter data). Now, if one is able to classify schizophrenia with above-chance accuracy from gray matter data, one cannot be sure which source of information within the data is picked up by the decoder: the actual neural representation of schizophrenia or the neural representation of gender or substance abuse? If one is interested in more than mere accurate *prediction* of schizophrenia, then this ambiguity due to confounding sources of information becomes problematic.

Importantly, as our definition suggests, what *is or is not* regarded as a confound is relative—it depends on whether the researchers deems it of (primary) interest or not. In the aforementioned hypothetical schizophrenia decoding study, for example, one may equally well define the severity of substance abuse as the to-be-decoded variable, in which the variable “schizophrenia-or-not” becomes the confounding variable. In other words, one researcher’s signal is another researcher’s confound. Regardless, if decoding analyses of neuroimaging data are affected by confounds, the data thus contain two types of information: the "true signal" (i.e., variance in the data related to the target, but unrelated to the confound) and the "confounded signal" (i.e., variance in the data related to the target that is also related to the confound; see Figure 1). In other words, representational ambiguity arises due to the presence of both true signal and confounded signal and, thus, controlling for confounds (by removing the confounded signal) provides a crucial methodological step forward in improving the interpretability of decoding analyses. In the following section, we will review the previously proposed methods to control for confounds.

**Figure 1.**
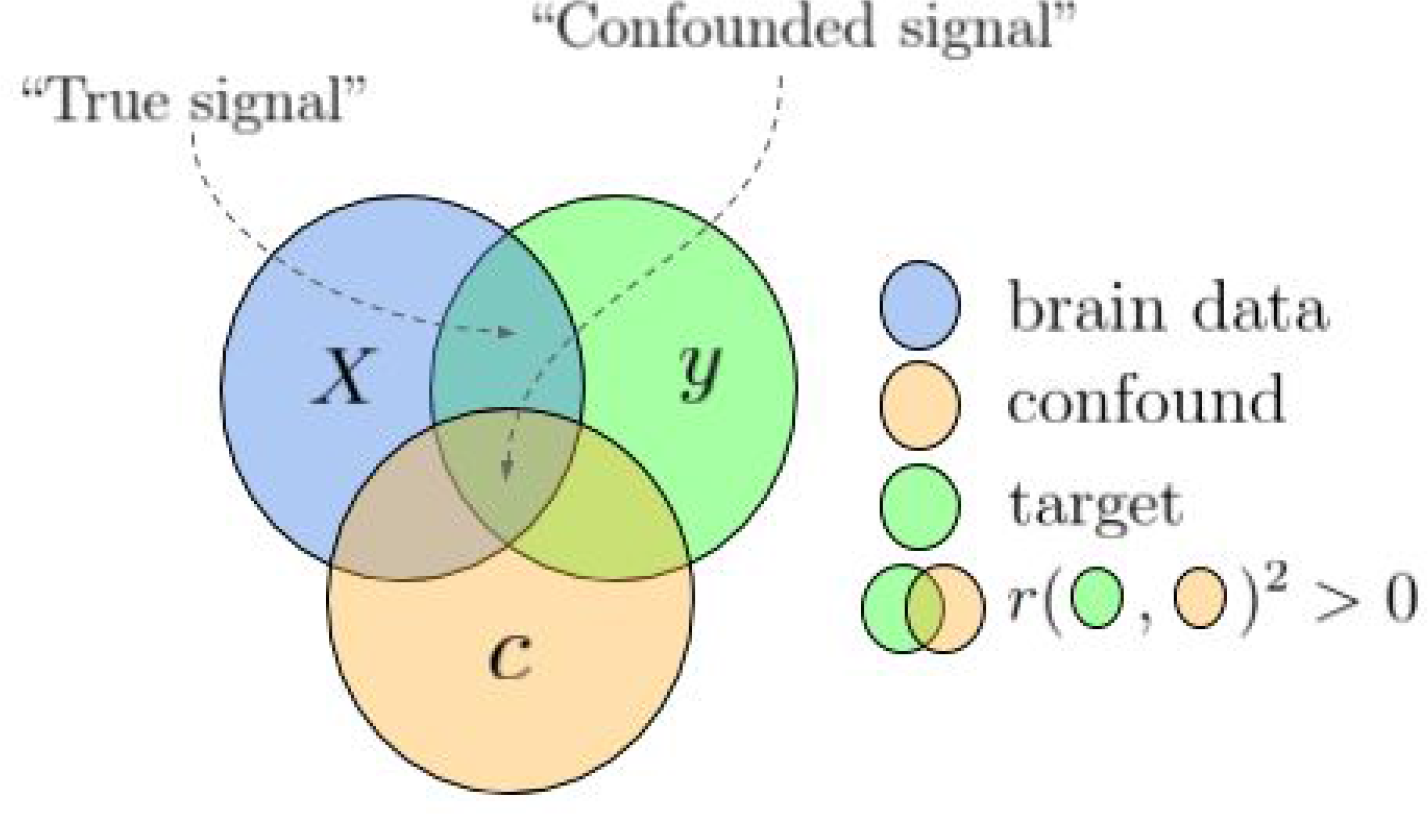
Visualization of how variance in brain data (*X*) can partitioned into “true signal” and “confounded signal”, depending on the correlation structure between the brain data (*X*), the confound (*C*), and the target (*y*). Overlapping circles indicate a non-zero (squared) correlation between the two variables.

### Methods for confound control

In decoding analyses, one aims to predict a certain target variable from patterns of neuroimaging data. In this section, we will review some common methods for controlling confounds in the context of decoding analyses, some of which are supplemented with a mathematical formalization; for consistency and readability, we define the notation we will use in Table 1.

**Table 1.** Notation.

| Symbol | Dims. | Description |
| --- | --- | --- |
| $N$ | | Number of samples (usually subjects or trials) |
| $K$ | | Number of neuroimaging features (e.g., voxels or sensors) |
| $P$ | | Number of confound variables (e.g., age, reaction time, or brain size) |
| $X$ | $N \times K$ | The neuroimaging patterns (often called the ‘data’ in the current article), where the subscript $i \in \{1, \dots, N\}$ refers to individual samples (rows), and the subscript $j \in \{1, \dots, K\}$ to individual features (columns) |
| $y$ | $N \times 1$ | The target variable (i.e., what is to be decoded) |
| $C$ | $N \times P$ | The confound variable(s) |
| $\beta$ | $K + 1$ | The parameters estimated in a general linear model (GLM) |
| $r_{Cy}$ | | Sample Pearson correlation coefficient(s) between $C$ and $y$ |
| $r_{y(X.C)}$ | | Sample semipartial Pearson correlation coefficient between $X$ and $y$, controlled for $C$ (i.e., $C$ is regressed out of $X$) |
| $p(r_{Cy})$ | | $p$-value of the sample Pearson correlation between $C$ and $y$ |

*Note:* format based on (Diedrichsen & Kriegeskorte, 2017). For the correlations, we assume that $P = 1$ and thus that the correlations in the table reduce to a scalar.

### A priori counterbalancing

Ideally, one would prevent confounding influences as much as possible before the acquisition of the neuroimaging data^3^. One common way (in both traditional ‘activation-based’ and decoding analyses) to prevent confounding is to make sure that potential confounding variables are *counterbalanced* in the experimental design (Görgen, Hebart, Allefeld, & Haynes, 2017). In the gender/brain size example described earlier, counterbalancing would entail ensuring that brain size does not differ significantly between men and women (i.e., given that men on average have larger brains than women, this would entail including only men with relatively small brains and women with relatively large brains)^4^.

Formally, in decoding analyses, a design is counterbalanced when the confound *C* and the target *y* are statistically independent. In practice, this often means that the sample is chosen so that there is no significant correlation coefficient between *C* and *y* (although this does not necessarily imply that *C* and *y* are actually independent). To illustrate the process of counterbalancing, let’s consider another hypothetical experiment: suppose one wants to set up an fMRI experiment in which the goal is to decode abstract object category (e.g. faces vs. houses) from the corresponding fMRI patterns (cf. Haxby et al., 2001), while controlling for the potential confounding influence of low-level or mid-level stimulus features, like luminance, spatial frequency, or texture (Long, Yu, & Konkle, 2017). Proper counterbalancing would entail making sure that the images used for this particular experiments have similar values for these low-level features across object categories (again see Görgen et al., 2017, for details). Thus, in this example, low-level stimulus features should be counterbalanced with respect to object category, such that above chance decoding of object category cannot be attributed to differences in low-level stimulus features (i.e., the confounds).

A priori counterbalancing of potential confounds is, however, not always feasible. For one, the exact measurement of a potentially confounding variable may be unknown until data acquisition. For example, brain size of a participant is only known after data collection. Similarly, Todd and colleagues (2013) found that their decoding analysis of rule representations was confounded by reaction times corresponding to the to-be-decoded trials. Another example of a “data-driven” confound is motion during data acquisition (important in, for example, decoding of clinical populations, like ADHD; Yu-Feng et al., 2007). In addition, counterbalancing confounds a priori may be challenging because of limited clinical populations, in which researchers do not have the luxury of maintaining a counterbalanced sample due to small sample sizes. Lastly, researchers may simply discover confounds after data acquisition.

Given that a priori counterbalancing is not possible or undesirable in many situations, it is paramount to explore the possibilities of controlling for confounding variables after data acquisition for the sake of model interpretability. From the neuroimaging literature, we identified several methods that aim to control for confounds in decoding analyses, which we discuss below in turn.

### Include confounds in model

One perhaps intuitive method to control for confounds in decoding analyses is to include (or, technically, column-wise concatenate) the confounds to the set of predictors (i.e., the neuroimaging data, *X*; see, e.g., Sepehrband et al., 2018). This intuition may stem from the analogous situation in univariate (activation-based) analyses of neuroimaging data, in which confounding variables are similarly controlled for by including them in the design-matrix together with the stimulus/task regressors. For example, in univariate analyses of functional MRI, movement of the participant is often controlled for by including motion estimates in the design matrix of first-level analyses (Johnstone et al., 2006); in EEG, some control for activity due to eye-movements by including activity measured by concurrent electro-oculography as covariates in the design-matrix (Parra, Spence, Gerson, & Sajda, 2005). Usually, the general linear model is then used to estimate each predictor’s influence on the neuroimaging data. Importantly, the parameter estimates are often interpreted as reflecting the unique contribution of each predictor variable, independent from the influence of the confound.

In the context of decoding, however, this arguably intuitive method to control for confounds by including them as predictors is problematic. This is because this method neglects the fact that these two types of analyses perform inference on different statistics: univariate (activation-based) analyses usually focus on parameter estimates (β), while decoding analyses focus on model performance (usually measured as explained variance, *R^2^*, or classification accuracy; Hebart & Baker, 2017). While including confounds in the model effectively controls for the *parameter values* of predictors-of-interest^5^, this method does not control the value for *model fit*. Model performance statistics (e.g., *R^2^*, classification accuracy, etc.) alone cannot disentangle the contribution of different sources of information as it only represents a single summary statistic of model fit (Ritchie, Kaplan, & Klein, 2017), which will only increase when adding more predictors to the model (especially if these predictors are correlated to the target, as is the case with confounds). One might, then, argue that additionally inspecting parameter values of decoding models may help in disambiguating different sources of information (Sepehrband et al., 2018). However, it has been shown that the weight and direction of those parameters cannot reliably be mapped to specific sources of information, i.e., as being task- or confound-related (e.g., features with large weights may be completely uncorrelated to the target variable; Haufe et al., 2014). As such, it does not make sense to include confounds to the set of predictors when the goal is to disambiguate the different sources of information in decoding analyses.

Recently, another approach similar to including confounds in the model has been proposed, which is based on the idea of a dose-response curve (Alizadeh, Jamalabadi, Schönauer, Leibold, & Gais, 2017). In this method, instead of adding the confound(s) to the model directly, the relative contribution of true and confounded signal is systematically controlled. The authors show that this approach is able to directly quantify the *unique* contribution of each source of information, thus effectively controlling for confounded signal. However, while sophisticated in its approach, this method only seems to work for categorical confounds, as it is difficult (if not impossible) to systematically vary the proportion of confound-related information when dealing with continuous confounds or when dealing with more than one confound.

### Control for confounds during spattern estimation

Another method that has been used by some decoding studies on functional MRI data deals with confounds in the initial procedure of estimating activity patterns of the to-be-decoded events (Woolgar, Golland, & Bode, 2014). In this method, an initial first-level (univariate) analysis models the fMRI time series (*s*) as a function of both predictors-of-interest (*X*) and the confounds (*C*), often using the GLM^6^:

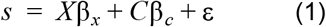

Then, *only* the estimated parameters (β^̂^, or normalized parameters, such as *t*-values or *z*-values) corresponding to the predictors-of-interest (β^̂^_*x*_) are used as activity estimates (i.e., the *X* used for predicting the target *y*) in the subsequent decoding analyses. This method thus takes advantage of the shared variance partitioning in the pattern estimation step to control for potential confounding influences. However, while elegant in principle, this method is not applicable in between-subject decoding studies (e.g. clinical decoding studies; e.g., van Waarde et al., 2014; Cuingnet et al., 2011), in which confounding variables are defined across subjects, or in electrophysiology studies, in which activity patterns do not have to be estimated in a first-level model^7^, thus limiting the applicability of this method.

### Post-hoc counterbalancing of confounds

When *a priori* counterbalancing is not possible, some have argued that *post-hoc* counterbalancing might control for the influence of confounds (Rao et al., 2017, p. 24, 38). In this method, given that there is some sample correlation between the target and confound (*r_Cy_* ≠ 0) in the entire dataset, one takes a subset of samples in which there is no empirical relation between the confound and the target anymore (e.g., when *r_Cy_* ≈ 0). In other words, post-hoc counterbalancing is a way to *decorrelate* the confound and the target through subsampling the data. Then, subsequent decoding analysis on the subsampled data can only capitalize on true signal, as there is no confounded signal anymore after subsampling (see Figure 2). While intuitive in principle, we are not aware whether this method has been evaluated before and whether it yields unbiased performance estimates.

**Figure 2.**
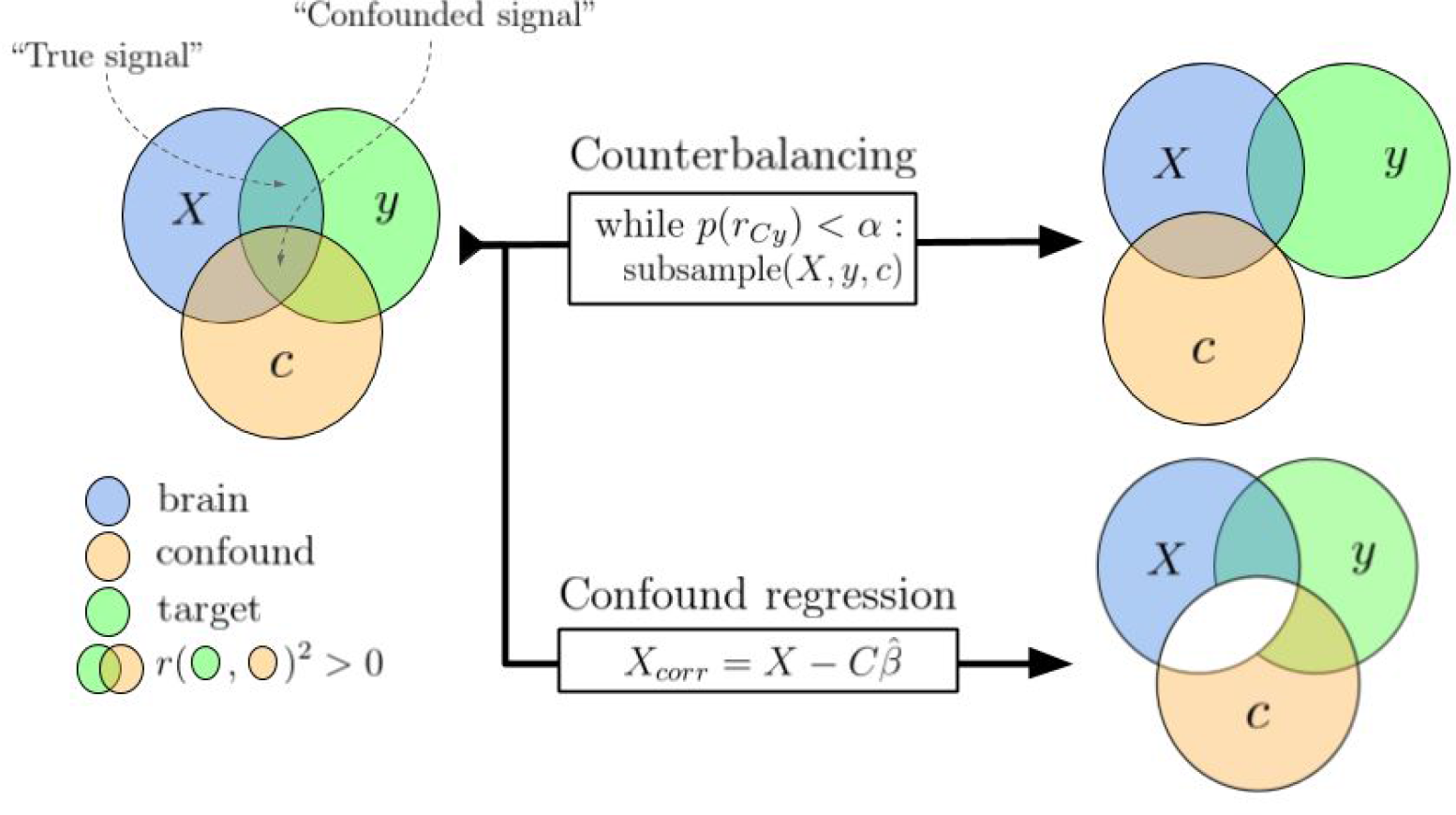
A schematic visualization how the main two confound control methods evaluated in this article deal with the “confounded signal”, making sure decoding models only capitalize on the “true signal”.

### Confound regression

The last and perhaps most common method to control for confounds is by removing the variance of the confound (i.e., the confounded signal) from the neuroimaging data directly (Abdulkadir, Ronneberger, Tabrizi, & Klöppel, 2014; Dukart, Schroeter, Mueller, Initiative, & Others, 2011; Kostro et al., 2014; Rao, Monteiro, Mourao-Miranda, & Alzheimer’s Disease Initiative, 2017; Todd et al., 2013) — a process we refer to as *confound regression* (also known as “image correction”; Rao et al., 2017). In this method, a (usually linear) regression model is fit on each feature in the neuroimaging data (i.e., a single voxel or sensor) with the confound(s) as predictor(s). Thus, each feature *j* in the neuroimaging data *X* is modelled as a linear function of the confounding variable(s), *C* :

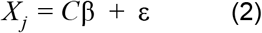

We can estimate the parameter(s) β̂_*j*_ for feature *X_j_* using, for example, ordinary least squares as follows (but for an example using a different model, see Abdulkadir et al., 2014):

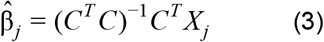

Then, to remove the variance of (or "regress out") the confound from the neuroimaging data, we can subtract the variance in the data associated with confound (*C*β̂_*j*_) from the original data:

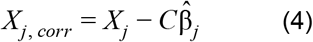

In which *X_j, corr_* represents the neuroimaging feature *X_j_* from which all variance of the confound is removed (including the variance shared with *y*, i.e., the confounded signal; see Figure 2). When subsequently applying a decoding analysis on this corrected data, one can be sure that the decoder is not capitalizing on signal that is correlated to the confound, which thus improves interpretability of the decoding analysis.

Confound regression has been applied in several studies in the context of decoding^8^. Todd and colleagues (2013) were, as far as the current authors are aware, the first to use this method to control for a confound (in their case, reaction time) that was shown to correlate with their target variable (rule A vs. rule B). Notably, they both regressed out reaction time from the first-level time series data (similar to the “Control for confounds during pattern-estimation” method) *and* regressed out reaction time from the trial-by-trial activity estimates (i.e., confound regression as described in this section). They showed that controlling for reaction time in this way completely eliminated the above-chance decoding performance in a substantial amount of voxels that was observed when not controlling for reaction time. Similarly, Kostro et al. (2014) observe a substantial drop in classification accuracy when controlling for scanner-site in the decoding analysis of Huntington’s disease, but only when scanner-site and disease status were actually correlated. Lastly, Rao and colleagues (2017) found that, in contrast to Kostro et al. and Todd et al., confound regression yielded similar (or slightly lower, but still significant) performance compared to the model without confound control, but it should be noted that this study used a regression model (instead of a classification model) and evaluated confound control in the specific situation when the training set is confounded, but the test-set is not^9^. In sum, while confound regression has been used before, it has yielded variable results, possibly due to slightly different approaches and differences in the correlation between the confounding variable and the target.

### Current study

In summary, multiple methods have been proposed to deal with confounds in decoding analyses. Often, these methods have specific assumptions about the nature or format of the data (like “A priori counterbalancing” and “Confound control during pattern estimation”), differ in their objective (e.g., *prediction* vs. *interpretation*, like in “Include confounds in the model”), or have yielded variable results (like “Confound regression”). Therefore, given that we are specifically interested in disambiguating decoding analyses, the current study evaluates the two methods that are applicable in most contexts: post-hoc counterbalancing and confound regression. In addition to these two methods, we propose a third method—a modified version of confound regression—which we show yields plausible, unbiased, and interpretable results.

To test whether these methods are able to effectively control for confounds and whether they yield unbiased results, we apply them both to empirical data and simulated data in which the ground truth with respect to the signal in the data (i.e., the proportion of true signal and confounded signal) is known. For our empirical data, we enact the previously mentioned hypothetical study in which participant gender is decoded from structural MRI data. We use a large dataset (*N* = 217) with structural MRI data and try to predict subjects’ gender (male/female) from the respective gray matter patterns while controlling for the confound of "brain size" using the aforementioned methods, which we compare to a baseline model in which confounds are not controlled for. Given the previously reported high correlations between brain size and gender (Barnes et al., 2010; Smith & Nichols, 2018), we expect that successfully controlling for brain size yields lower decoding performance than using uncorrected data, but not below chance level. Note that higher decoding performance after controlling for confounds is theoretically possible when the correlation between the confound and the target is sufficiently low to cause suppressor effects (see Figure 1 in Haufe et al., 2014). However, because our confound, brain size, is known to correlate strongly with our target, gender (approx. *r* = 0.63; Smith & Nichols, 2018), classical suppression effects are unlikely and thus we expect lower model performance after controlling for brain size.

However, our results indicate that both counterbalancing and confound regression lead to unexpected results: counterbalancing fails to reduce model performance while confound regression consistently yields very low model performance up to the point of significant below-chance accuracy. In subsequent simulations, we show that both methods lead to *biased* results: counterbalancing yields inflated model performance (i.e., positive bias) because subsampling selectively selects a subset of samples in which features correlate more strongly with the target variable, suggesting (indirect) circularity in the analysis (Kriegeskorte, Simmons, Bellgowan, & Baker, 2009). Furthermore, our simulations show that negative bias (including significant below-chance classification) after confound regression on the entire dataset is due to reducing the signal below what is expected by chance (Jamalabadi, Alizadeh, Schönauer, et al., 2016), which we show is related to and can be predicted by the standard deviation of the empirical distribution of correlations between the features in the data and the target. We propose a minor but crucial addition to the confound regression procedure, in which we cross-validate the confound regression models (which we call “Foldwise Confound Regression”), which solves the below-chance accuracy issue and yields plausible model performance in both our empirical and simulated data.

## METHODS

### Data

For the empirical analyses, we used voxel-based morphometry (VBM) data based on T1-weighted scans and tract-based spatial statistics (TBSS) data based on diffusion tensor images from 217 participants (122 women, 95 men), acquired with a Philips Achieva 3T MRI-scanner and a 32-channel head coil at the Spinoza Centre for Neuroimaging (Amsterdam, The Netherlands).

#### VBM acquisition & analysis

The T1-weighted scans with a voxel size of 1.0 × 1.0 × 1.0 mm were acquired using 3D fast field echo (TR: 8.1 ms, TE: 3.7 ms, flip angle: 8°, FOV: 240 × 188 mm, 220 slices). We used "FSL-VBM" protocol (Douaud et al., 2007) from the FSL software package (version 5.0.9; (Smith et al., 2004) using default and recommended parameters (including non-linear registration to standard space). The resulting VBM-maps were spatially smoothed using 3 millimeter (FWHM) gaussian kernel. Subsequently, we organized the data in the standard pattern-analysis format of a 2D (*N* × *K*) array of shape 217 (subjects) × 412473 (non-zero voxels).

#### TBSS acquisition & analysis

Diffusion tensor images with a voxel size of 2.0 × 2.0 × 2.0 mm were acquired using a spin-echo echo-planar imaging (SE-EPI) protocol (TR: 7476 ms, TE: 86 ms, flip angle: 90°, FOV: 224 × 224 mm, 60 slices), which acquired a single b=0 (non-diffusion-weighted) image and 32 (diffusion-weighted) b=1000 images. All volumes were corrected for eddy-currents and motion (using the fsl command ‘eddy_correct’) and the non-diffusion-weighted image was skullstripped (using FSL-BET with the fractional intensity threshold set to 0.3) to create a mask that was subsequently used in the fractional anisotropy (FA) estimation. The FA-images resulting from the diffusion tensor fitting procedure were subsequently processed by FSL’s tract-based spatial statistics (TBSS) pipeline (Smith et al., 2006), in which we used the pipeline’s recommended parameters (i.e., non-linear registration to FSL’s 1 millimeter FA image, construction of mean FA-image and skeletonized mean FA-image based on the data from all subjects, and a threshold of 0.2 for the skeletonized FA-mask). Subsequently, we organized the resulting skeletonized FA-maps into a 2D (*N* × *K*) array of shape 217 (subjects) × 128340 (non-zero voxels).

#### Brain size estimation

To calculate the values for our confound, global brain size, we calculated for each subject separately the total number of nonzero voxels in the gray matter and white matter map resulting from the segmentation step in the FSL-VBM pipeline (using FSL’s segmentation algorithm "FAST"; Zhang, Brady, & Smith, 2001). The number of non-zero voxels from the gray matter map was used as the confound for the VBM-based analyses and the number of non-zero voxels from the white matter map was used as the confound for the TBSS-based analyses. Note that brain size estimated from total white matter volume and total gray matter volume correlated strongly, *r*(216) = 0.93, *p* < 0.001.

#### Data and code availability

In the Github repository corresponding to this article (https://github.com/lukassnoek/MVCA), we included a script to download the data (the 4D VBM and TBSS nifti-images as well as the non-zero 2D samples × features arrays). The repository also contains detailed Jupyter notebooks with the annotated empirical analyses and simulations reported in this article.

### Decoding pipeline

All empirical analyses and simulations use a common decoding pipeline, which is implemented using functionality from the *scikit-learn* Python package for machine learning (Abraham et al., 2014; Pedregosa et al., 2011). This pipeline includes univariate feature selection (based on a prespecified amount of voxels with highest univariate difference in terms of the ANOVA *F*-statistic), feature-scaling (ensuring zero mean and unit standard deviation for each feature), and a support vector classifier (SVC) with a linear kernel, fixed regularization parameter (*C* = 1), and sample weights set to be inversely proportional to class frequency (to account for class imbalance). In our empirical analyses, we evaluate model performance for different amounts of voxels (as selected by the univariate feature selection). We report model performance as the *F*_1_ score, which is insensitive to class imbalance (which, in addition to adjusted sample weights, prevents the classifier to learn the relative probabilities of target classes instead of representative information in the features). The F_1_ score has an expected chance level of 0.5. Statistical significance was calculated using non-parametric permutation tests as implemented in scikit-learn with 1000 permutations (Ojala & Garriga, 2010).

### Evaluated methods for confound control

#### Counterbalancing

We implement post-hoc counterbalancing in two steps. First, to quantify the relation between the confound and the target in our dataset, we estimate the point-biserial correlation coefficient between the confound, *C* (brain size), and the target, *y* (gender) across the entire dataset (all samples *i* = 1, …, *N*). Because of both sampling noise and measurement noise, sample correlation coefficients vary around the population correlation coefficient and are thus improbable to be 0 *exactly*^10^. Therefore, in the next step, we subsample the data until the correlation coefficient between *C* significance threshold α :

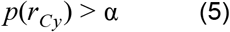

In our analyses, we use an α of 0.1.^11^

Then, given that the subsampled dataset is counterbalanced with respect to the confound, a random stratified K-fold cross-validation scheme is repeatedly initialized until a scheme is found in which *all* splits are counterbalanced as well (cf. Görgen et al., 2017). This particular counterbalanced cross-validation scheme is subsequently used to cross-validate the MVPA pipeline. We implemented this counterbalancing method as a *scikit-learn*-style cross-validator class, available from the aforementioned Github repository (in the counterbalance.py module).

#### Confound regression

In our empirical analyses and simulations, we tested two different versions of confound regression, which we call "whole-dataset confound regression" (WDCR) and "foldwise confound regression" (FwCR). In WDCR, we regress out the confounds from the predictors *from the entire dataset at once*, i.e., before entering the iterative cross-validated MVPA pipeline (the approach taken by Abdulkadir et al., 2014; Dubois, Galdi, Han, Paul, & Adolphs, 2017; Kostro et al., 2014; Todd et al., 2013). Note that we can do this for all *K* voxels at once using the closed-form OLS solution, in which we first estimate the parameters β̂_*C*_ :

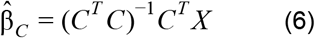

In which *C* is an *N* × 2 array in which the first column contains an intercept and the second column contains the confound brain size. Accordingly,β̂_*C*_ is an 2 × *K* array. We then remove the variance associated with the confound from our neuroimaging data as follows:

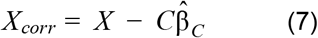

Now, *Xcorr* is an array with the same shape as the original *X* array, but with the variance related to the confound *C* removed (i.e., *X* is residualized with regard to *C*).

In our proposed cross-validated version of confound regression (which was mentioned but not evaluated by Rao et al., 2017, p. 25), "FwCR", we similarly regress out the confounds from the neuroimaging data, yet instead of estimating β̂*_C_* on the entire dataset, we estimate this *within each fold of training data* (*X_train_*):

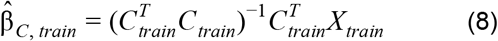

And we subsequently use these parameters (β^̂^_*C*, *train*_) to remove the variance related to the confound from both the train-set (*X_train_* and *C_train_*):

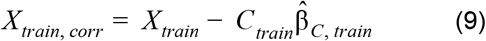

and the test-set (*X_test_* and *C_test_*):

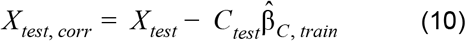

Thus, essentially, FwCR represents the cross-validated version of WDCR. We implemented these confound regression techniques as a *scikit-learn* compatible transformer object, available in the open-source *skbold* Python package (Snoek, 2017).

#### Simulations

In addition to the empirical evaluation of counterbalancing and confound regression in the gender decoding example, we ran three simulations: one generic simulation to evaluate the three confound control methods on synthetic data (“generic simulation”), one simulation to specifically investigate positive bias observed after counterbalancing (“counterbalancing follow-up simulation”), and one simulation to specifically investigate the negative bias after WDCR and to demonstrate FwCR solves this negative bias (“WDCR/FwCR follow-up simulation”). We discuss the implementation of these simulations below.

#### Generic simulation

In this simulation, we evaluate how the three methods for confound control behave on data with a prespecified correlation between the confound and the target, *r_Cy_*, and different amounts of "confounded signal" (i.e., the explained variance in *y* driven by shared variance between *X* and *C*). These simulations allow us to have full control over (and knowledge of) the influence of the signal and confound in the data, and thereby help diagnosing the issues associated with counterbalancing and confound control (which are investigated in detail in the method-specific simulations).

Specifically, in the generic simulation, we generate hypothetical data sets holding the correlation coefficient between *C* and *y* constant, while varying the amount of true signal and confounded signal. We operationalize true signal as the the squared semipartial Pearson correlation between *y* and each feature in *X*, controlled for *C*. As such, we will refer to this term as *signal R^2^*:

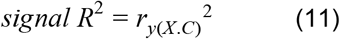

Similarly, we operationalize the confounded signal as the shared explained variance of *y* by each feature of *X* and *C*. This term, which we will refer to as *confound R^2^*, is calculated as follows:

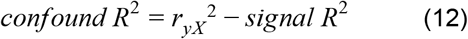

In the simulations reported and shown in the main article, we use *r_Cy_* = 0.65, which corresponds to the observed correlation between brain size and gender in our dataset. To create data with this prespecified structure, within our simulations we generated (1) a data-matrix *X* of shape *N* × *K*, (2) a target variable *y* of shape *N* × 1, and (3) a confound variable *C* of shape *N* × *P*. For the simulations reported in this manuscript, we generate data with *N* = 200, *K* = 5, and *P* = 1 (i.e., a single confound variable). We generate *y* as a categorical variable with binary values, *y* ∈ {0, 1}, with equal class probabilities (i.e., 50%), given that most decoding studies focus on binary classification. We generate *C* as a continuous random variable drawn from a standard normal distribution. We generate each feature *X_j_* as a linear combination of *y* and *C* plus Gaussian noise. Thus, for each predictor *j* = 1, …, *K* in *X_j_* :

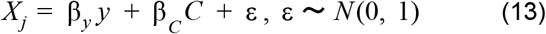

in which β_*y*_ represents the weight given to *y* and β_*C*_ represents the weight given to *C* in the generation of the feature *X_j_*, and *N* (*a*, *b*) is the normal distribution with mean *a* and standard deviation *b*. Then, we partition the explained variance of *y* into signal *R^2^* and confound *R^2^*. If one or both of these values are off by more than 0.01 from the desired values, the generative parameters β_y_ and β_C_ are adjusted after which *X_j_* is generated again, which is iterated until the data contain the desired "true signal" and "confounded signal". Here, we evaluate the different methods for confound control for two different values for signal *R^2^* (0.004, representing plausible null data^12^, and 0.1, representing a plausible true effect) and a range of confound *R^2^* values (in steps of 0.05: 0.00, 0.05, 0.10, …, 0.35). This simulation is iterated 10 times (with different partitions of the folds) for robustness. Importantly, the specific scenario in which confound *R*^2^ equals 0, which represents data which quantitatively does not contain any confounded signal (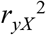 = *Signal R*^2^), will serve as “reference model performance” to which we can compare the efficacy and bias of the confound control methods.

After the data have been generated, a baseline model (no confound control) and the three methods outlined above (counterbalancing, WDCR, and FwCR) are applied to the simulated data using the standard pipeline described in the “Decoding pipeline” section (but without univariate feature selection) and compared to the “reference performance”.

#### Counterbalancing follow-up simulation

To further investigate the positive bias after counterbalancing, we simulate a multivariate normal dataset with three variables, which reflect our data (*X*), target (*y*), and confound (*C*), with 1000 samples (*N*) and a single feature (*K* = 1). We iterate the process 1000 times and subsequently use the dataset which yields the largest (positive) difference between model performance after counterbalancing versus no confound control; in other words, we use the dataset in which the counterbalancing issue is most apparent. While not necessarily representative of typical (neuroimaging) datasets, this process allows us to clearly explain and visualize what causes the positive bias after counterbalancing the data.

To generate data from a multivariate normal distribution, we specified the variance-covariance matrix to have unit variance for all variables, so that covariances can be interpreted as correlations. The covariances in the matrix were generated as pairwise correlations (*r_yX_*, *r_CY_*, *r_CX_*), each sampled from a uniform distribution with range [− 0.65, 0.65]. We generate data using such pre-specified correlation structure because the relative improvement in model performance after counterbalancing does not appear to occur when generating completely random (normal) data. Moreover, we restrict the range of the uniform distribution from which the pairwise correlations are drawn to [− 0.65, 0.65] because a larger range may result in the covariance matrix not being positive-semidefinite. After generating the three variables, we binarize the target variable (*y*) using a mean-split (*y* = 0 if *y* < *y*̄, *y* = 1 otherwise) to frame the analysis as a classification problem rather than a regression problem.

We then subsample the selected dataset using on our post-hoc counterbalancing algorithm and subsequently run the decoding pipeline (without univariate feature selection) on the subsampled (“retained”) data in a 10-fold cross-validation scheme. Notably, we cross-validate our fitted pipeline not only to the left-out retained data, but also to the data that did not survive the subsampling procedure (the “rejected” data; see Figure 3). Across the 10 folds, we keep track of two statistics of both the retained and rejected samples: (1) whether the samples were predicted correctly and (2) the signed distance to the decision boundary. Negative distances in binary classification (in simple binary classification with *y* ∈ {0, 1}) reflect a prediction of the sample as *y* = 0, while positive distances reflect a prediction of the sample as *y* = 1. Here, however, we want to count the distance of samples that are on the “incorrect” side of the decision boundary as *negative* distances, while counting the distance of samples that are on the “correct” side of the decision boundary as *positive* distances. To this end, we use a “re-coded” version of the target variable (*y*^*^ =− 1 if *y* = 0, *y*^*^ = 1 otherwise) and multiply it with the distance. As such, we calculate the signed distance from the decision boundary (δ_*i*_) for any sample *i* as:

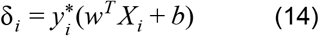

in which *w* refers to the feature weights (coefficients) and *b* refers to the intercept term. Then, differences in these two statistics (proportion correctly classified and distance to boundary) between the retained and rejected samples allows us to evaluate if counterbalancing yields unbiased estimates (i.e., better cross-validated model performance on the retained data than on the rejected data would confirm positive bias, as it indicates that subsampling tends to reject hard-to-classify samples). We apply this analysis similarly to the empirical data (separately for the different values of *K*) to show that the effect of counterbalancing as demonstrated using simulated data also occurs in the empirical data.

**Figure 3.**
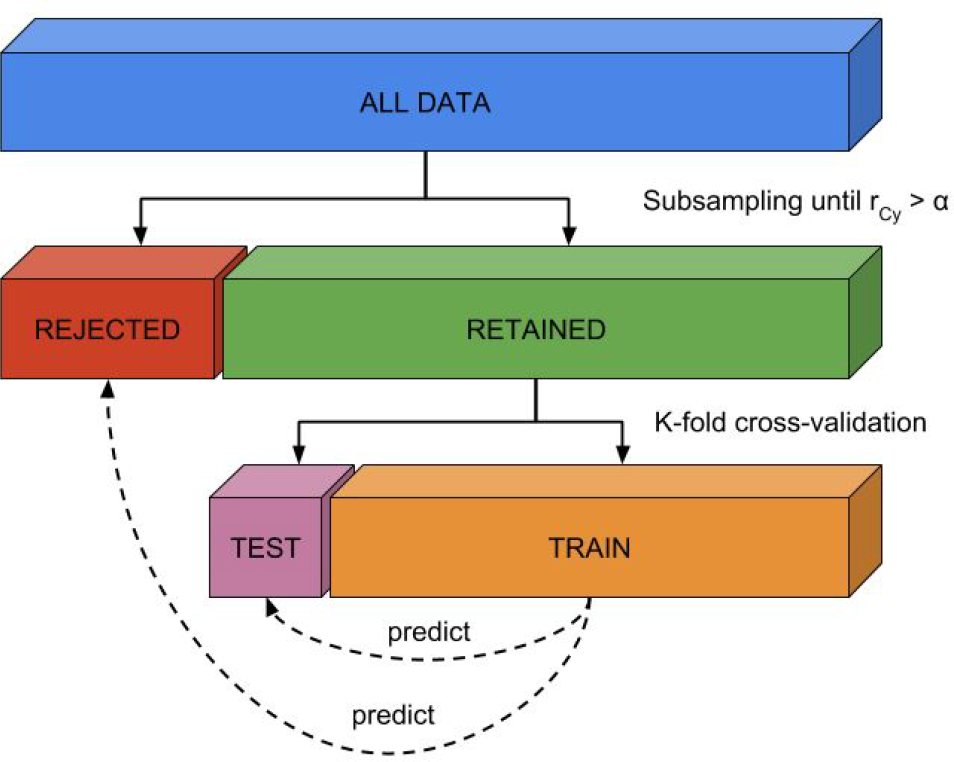
Visualization of method to evaluate whether counterbalancing yields unbiased cross-validated model performance estimates.

#### WDCR/FwCR follow-up simulation

To demonstrate the problem with WDCR (and to demonstrate that FwCR solves this), we perform two follow-up simulations: the first follow-up simulation shows that below-chance accuracy depends on the distribution of feature-target correlations (*r_yX_*; see Jamalabadi et al., 2016, for a similar argument) and the second follow-up simulation to show that WDCR artificially narrows this distribution, leading to below chance accuracy, which is exacerbated by increasing number of features (*K*) and higher correlations between the target and confound (*r_Cy_*). In the first simulation, we simulate random null-data (drawn from a standard normal distribution) with 100 samples (*N*) and 200 features (*K*), as well as a binary target feature (*y* ∈ {0, 1}). We then calculate the cross-validated accuracy using the standard pipeline (without univariate feature selection) described in the “Decoding pipeline” section; we iterate this process 500 times. Then, we show that the variance of the cross-validated accuracy is accurately predicted by the standard deviation (i.e., “width”) of the distribution of correlations between the features and the target (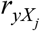 with *j* = 1, …, *K*), which we will denote by *sd*(*r_yX_*). Importantly, we show that below-chance accuracy likely occurs when the standard deviation of the feature-target correlation distribution is lower than the standard deviation of the sampling distribution of the Pearson correlation coefficient parameterized with the same number of samples (*N* = 200) and the same effect (i.e., ρ = 0, because we simulated random null-data). The sampling distribution of the Pearson correlation coefficient is described by Kendall & Stuart (1973). When as follows:

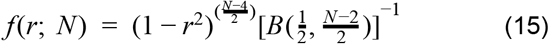

where *B*(*a*, *b*) represents the Beta-function.

Then, in a second simulation, we similarly simulate null-data as in the previous simulation, but now we also generate a continuous confound (*C*) with a variable correlation with the target (*r_Cy_* ∈ {0.0, 0.1, 0.2, …, 0.9, 1.0}). Before subjecting the data to the decoding pipeline, we regress out the confound from the data (i.e., WDCR). We do this for different amount of features (*K* ∈ {1, 5, 10, 50, 100, 500, 1000}). Then, we apply FwCR on the simulated data as well for comparison.

## RESULTS

### Influence of brain size

Before evaluating the different methods for confound control, we determined whether brain size is truly a confound given our proposed definition ("a variable that is not of primary interest, correlates with the target and is encoded in the neuroimaging data"). We evaluated the relationship between the target and the confound in two ways. First, we calculated the (point-biserial) correlation between gender and brain size, which was significant for both the estimation based on white matter, *r*(216) = .645, *p* < 0.001, and the estimation based on grey matter, *r*(216) = .588, *p* < 0.001, corroborating the findings by Smith & Nichols (2018). Second, as recommended by Görgen et al. (2017), who argue that the potential influence of confounds can be discovered by running a classification analysis using the confound as the (single) feature predicting the target, we ran our decoding pipeline (without univariate feature selection) using brain size as a single feature to predict gender. This analysis yielded a mean classification performance (*F*_1_ score) of 0.78 (*SD* = .10) when using brain size estimated from white matter and 0.81 (*SD* = .9) when using brain size estimated from gray matter, which are both significant with *p* < 0.001 (see Figure 4A).

**Figure 4.**
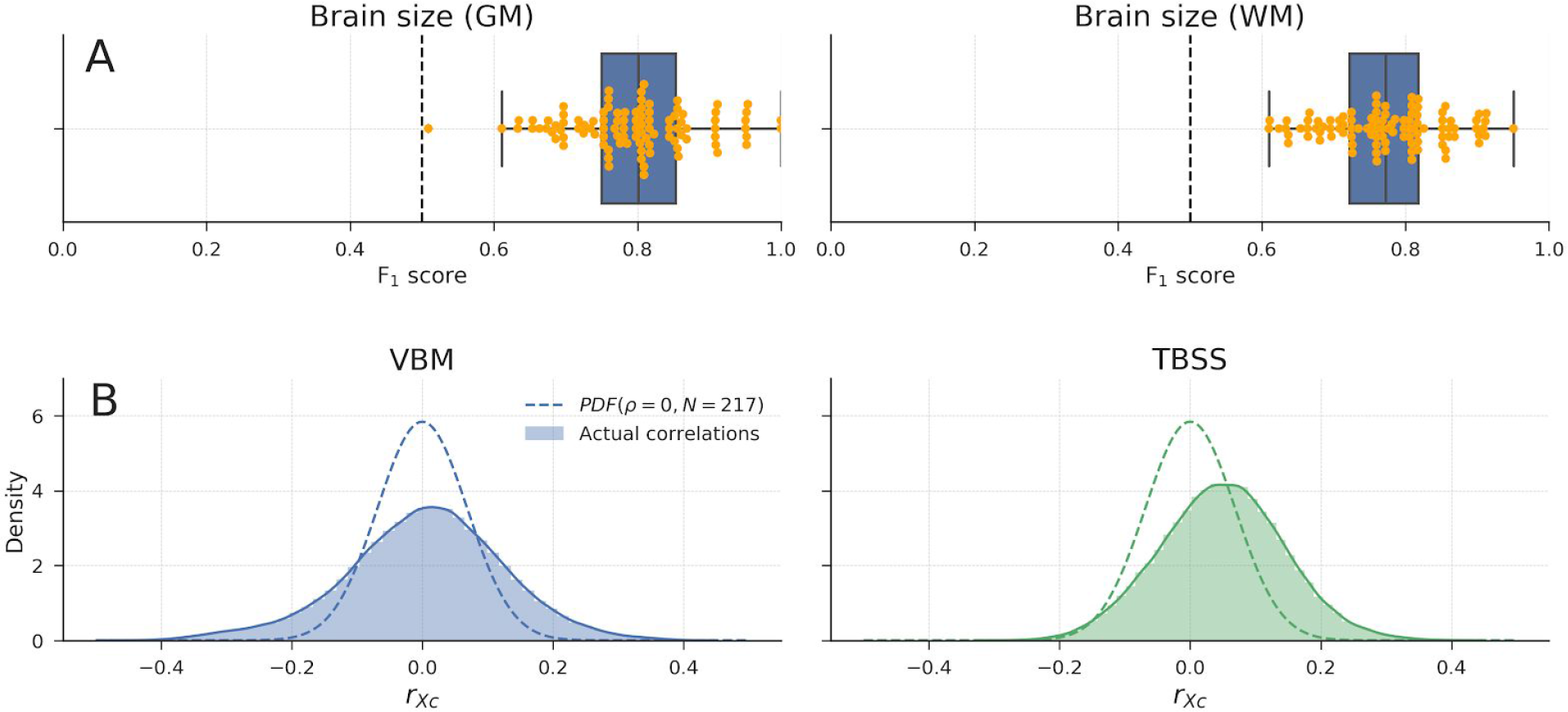
A) Model performance when using brain size to predict gender for both brain-size estimated from grey matter (left) and from white matter (right). Points in yellow depict individual F_1_ scores per fold in the 10-fold cross-validation scheme. Whiskers of the box plot are 1.5x the interquartile range. B) Distributions of observed correlations between brain size and voxels (*r_XC_*), overlayed with the analytic sampling distribution of correlation coefficients when ρ = 0 and *N* = 217, for both the VBM data (left) and TBSS data (right).

To estimate whether brain size is encoded in the neuroimaging data, we compared the distribution of bivariate correlation coefficients (of each voxel with brain size) with the sampling distribution of correlation coefficients when ρ = 0 and *N* = 217 (see section “WDCR/FwCR follow-up simulation” for details). Under the null hypothesis that there are no correlations between brain size and voxel intensities, each individual correlation coefficient between a voxel and the confound can be regarded as an independent sample with *N* = 217 (ignoring correlations between voxels for simplicity). Because *K* is very large for both the VBM and TBSS data, the empirical distribution of correlation coefficients should, under the null hypothesis, approach the analytic distribution of correlation coefficients parametrized by *N* = 217 and ρ = 0. Contrarily, the density plots in Figure 4B clearly show that the observed correlation coefficients distribution do not follow the sampling distribution (with both an increase in variance and a shift of the mode), indicating that at least some of the correlation coefficients between voxel intensities and brain size are extremely unlikely under the null hypothesis.

### Baseline model: no confound control

In our baseline model on the empirical data, for different amounts of voxels, we predicted gender from structural MRI data (VBM and TBSS) without controlling for brain size (see Figure 5). The results show significant above chance performance of the MVPA pipeline based on both the VBM-data and the TBSS-data. All performance scores averaged across folds were significant (*p* < 0.001).

**Figure 5.**
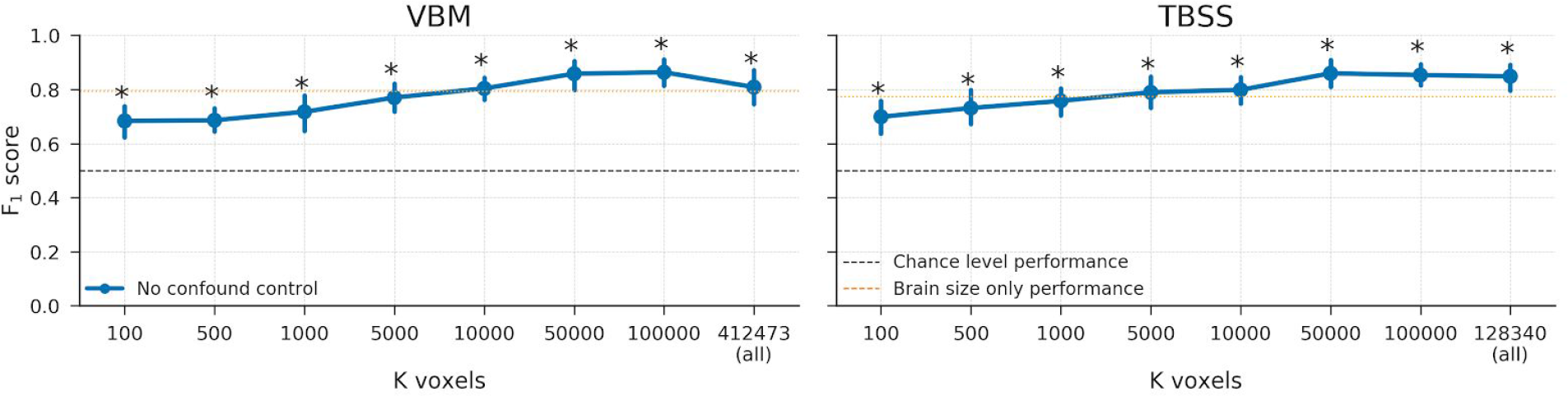
Baseline scores using the VBM (left) and TBSS (right) data without any confound control. Scores reflect the average F_1_ score across 10 folds; error bars reflect 95% confidence intervals across 1000 bootstraps. The dashed black line reflect theoretical chance-level performance and the dashed orange line reflects the average model performance when only brain size is used as a predictor for reference; * indicates significant performance above chance with *p* < 0.001.

These above-chance baseline performance estimates replicate previous studies on gender decoding from MRI data (Del Giudice et al., 2016; Rosenblatt, 2016; Sepehrband et al., 2018) and will serve as a baseline estimate of model performance to which the confound control methods will be compared. In the next three subsections, we test the three discussed methods to control for confounds: post-hoc counterbalancing, whole-dataset confound regression (WDCR), and foldwise confound regression (FwCR).

### Counterbalancing

#### Empirical results

In order to decorrelate brain size and gender (i.e., *r_Cy_* > 0.1), our subsampling algorithm selected 117 samples in the VBM data (i.e., a reduction in samples of 46.1%) and 131 samples in the TBSS data (i.e., a reduction of 39.6%). The model performance for different values of *K* (number of voxels) are shown in Figure 6. Contrary to our expectations, the predictive accuracy of our decoding pipeline is, for a substantial part of the results (especially in the VBM data), *higher* after counterbalancing than before counterbalancing. This is particularly surprising in light of the large reductions in sample size, which normally results in a hit in power and thus lower model performance.

**Figure 6.**
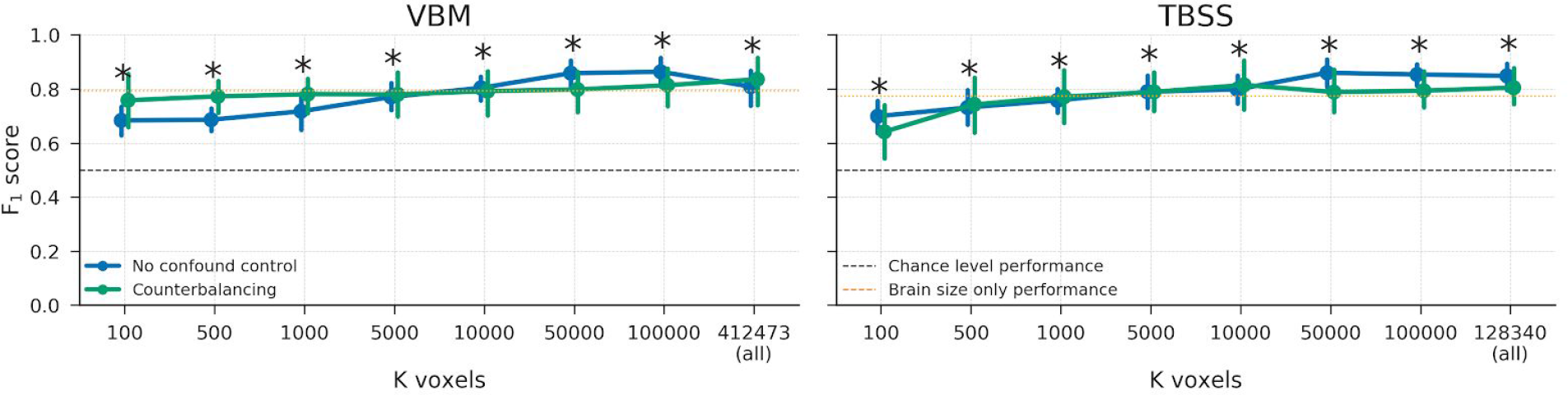
Model performance after counterbalancing (green) versus the baseline performance (blue) for both the VBM (left) and TBSS (right) data. Performance reflects the average F_1_ score across 10 folds; error bars reflect 95% confidence intervals across 1000 bootstraps. The dashed black line reflect theoretical chance-level performance (0.5) and the dashed orange line reflects the average model performance when only brain size is used as a predictor; * indicates significant performance (*p* & 0.001) above chance.

One could argue that this increase in model performance after counterbalancing can be explained by the possibility that the subsampling and counterbalancing just leads to the selection of different features using univariate feature selection compared to the baseline model. In other words, the increase in model performance may be caused by the feature selection function to select “better” voxels (i.e., containing better or more “robust” signal), which lead to higher model performance regardless of the reduction in sample size. However, this does not explain the similar scores for counterbalancing and the baseline model when using all voxels (the data points at *K voxels* = … (*all*) in Figure 6), as it is expected that successful confound control would yield a *lower* model performance given the high correlation between the target and the confound and the confound and the neuroimaging data (see “Influence of brain size” section).

Another possibility for the relative increase in performance of the model based on the counterbalanced data versus the baseline model is that counterbalancing increases the signal in the data. Indeed, counterbalancing appears to increase the (absolute) correlations between the data and the target (*r_yX_*), which is visualized in Figure 7, suggesting an increase in signal.

**Figure 7.**
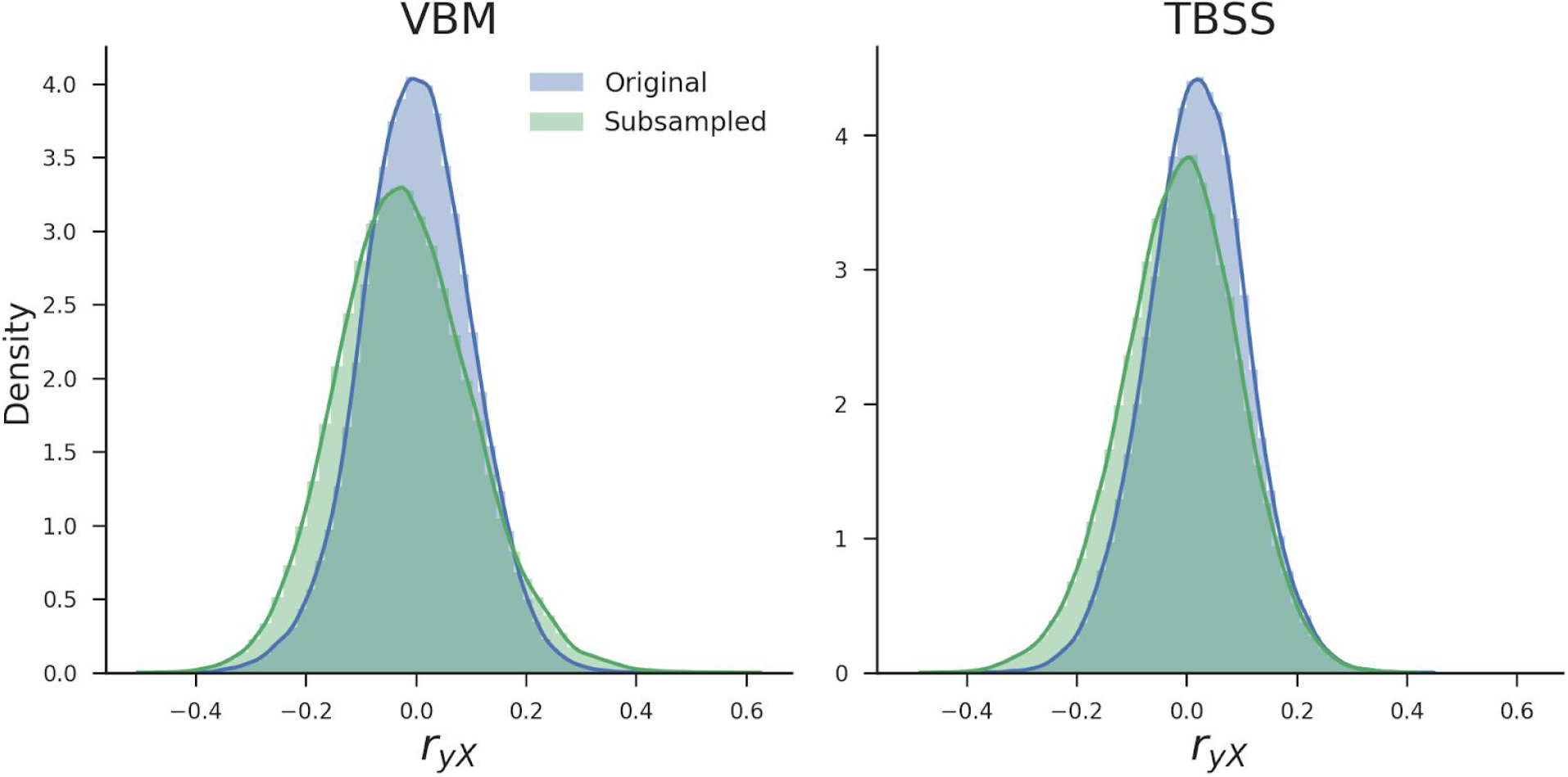
Density plots of the correlations between the target and voxels across all voxels before (blue) and after (green) subsampling for both the VBM and TBSS data

As can be seen in Figure 6 and Figure 7, counterbalancing increases the correlations between the target and neuroimaging data, which goes against the intuition that removing the influence of a confound that is highly correlated to the target will reduce decoding performance. To further investigate these issues, we first replicate this effect of counterbalancing on simulated data in our “generic simulation” section and then elucidate the mechanism behind this phenomenon in the “counterbalancing follow-up simulation”.

#### Generic simulation

In the generic simulation, we simulated data in which we varied the strength of *confound R^2^* and *signal R^2^*, after which we applied the three confound control methods to the data. The results from this generic simulation shows that counterbalancing maintains chance-level model performance when there is almost no signal (i.e., signal *R^2^* = 0.004; Figure 8, left graph, green line). However, when there is some signal (i.e., signal *R^2^* = 0.1; Figure 8, right graph), we see that counterbalancing yields similar or even higher scores than the baseline model, replicating the effects observed in the empirical analyses.

**Figure 8.**
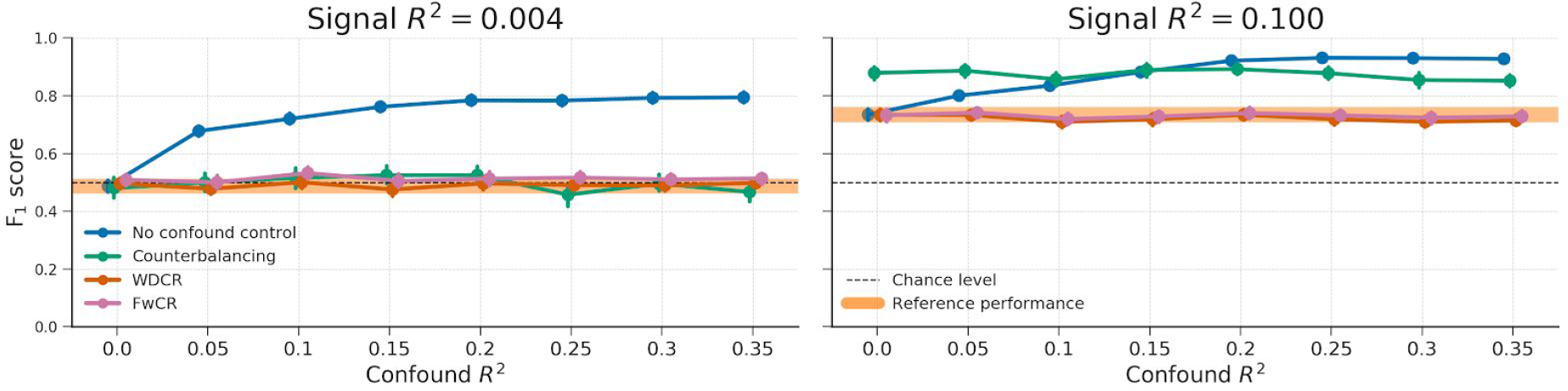
Results from the different confound control methods on simulated data without any experimental effect (signal *R^2^* = 0.004; left graph) and with some experimental effect (signal *R^2^* = 0.1; right graph) for different values of confound *R^2^*. The orange line represents the average performance (± 1 *SD*) when confound *R^2^* = 0, which serves as a “reference performance” for when there is no confounded signal in the data. For both graphs, the correlation between the target and the confound, *r_yC_*, is fixed at 0.65. The results from the WDCR and FwCR methods are explained later in the paper.

As is apparent from Figure 8, the counterbalanced data seems to yield better performance than the baseline model only for relatively low confound *R^2^* values (confound *R^2^* < 0.15). As suggested by our findings in the empirical data (see Figure 7), we hypothesized that the observed improvement in model performance after counterbalancing is caused by the increase in correlations between the target and neuroimaging data. In support of this hypothesis, we find that the increase in correlations after counterbalancing (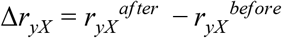) is indeed strongly correlated to the difference in performance between the counterbalancing model and the baseline model, *r*(79 = .922, *p* < 0.001 (see Figure 9), which seems to be inversely related to confound *R*^2^ (color-coded in Figure 9).

**Figure 9.**
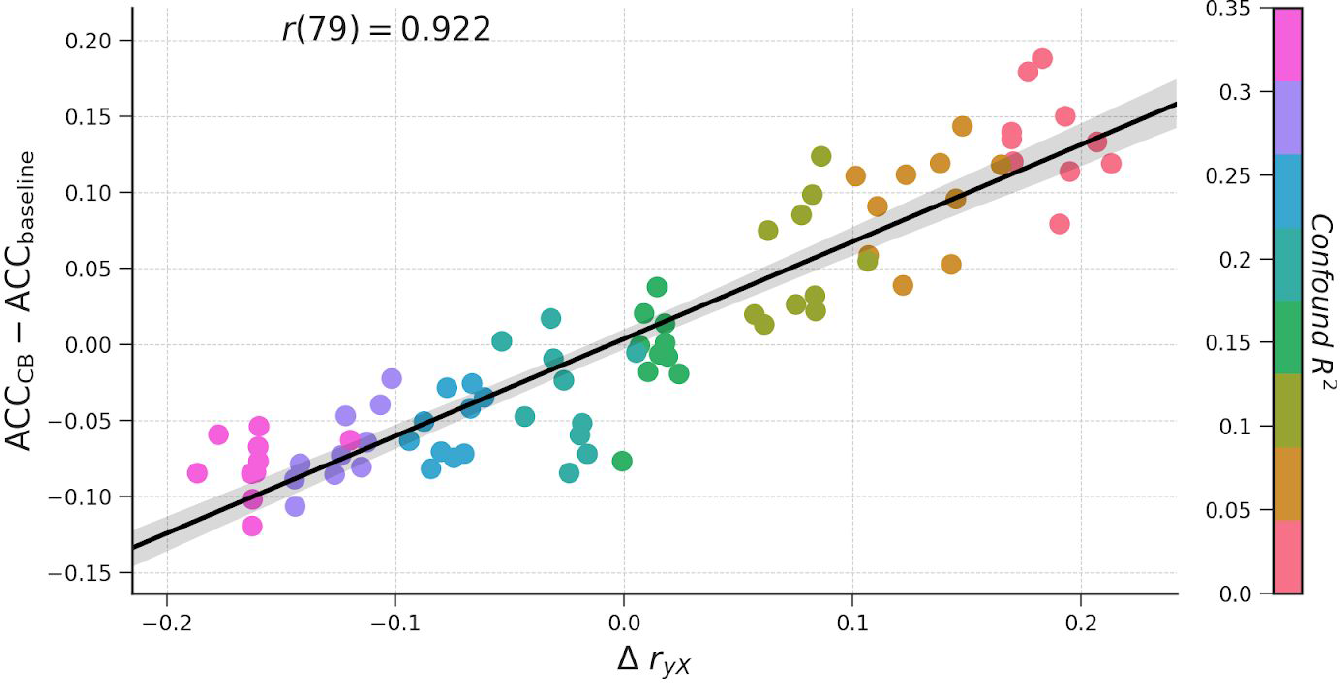
The relationship between the increase in correlations between target and data (*r_yX_*) after subsampling, confound *R^2^*, difference in model performance (here: accuracy) between the counterbalance model and baseline model (ACC_CB_ - ACC_baseline_).

While this relationship in Figure 9 might be statistically interesting, it does not tell us *why* counterbalancing tends to increase the correlations between neuroimaging data and target and even outperforms the baseline model when confound *R^2^* is low and there is signal present. More importantly, it does not tell us whether the counterbalancing procedure uncovers signal that is truly related to the target—in which case the confound acted as a suppressor before counterbalancing—or inflates performance estimates and thereby introduces positive bias. Therefore, in the next section we report and discuss results from a follow-up simulation that intuitively shows why counterbalancing leads to an increase in performance, and that this increase is in fact a positive bias.

#### Counterbalancing follow-up simulation

In this follow-up simulation, we aim to visualize the scenario in which counterbalancing leads to a clearly better performance than in case of no confound control. As such, we iteratively generate data with a correlation structure that we know leads to a large difference between counterbalancing and the baseline model (i.e., data with a low confound *R^2^*) and choose the variables (*X*, *y*, *C*) that yielded the largest difference for our visualization (see the “Counterbalancing follow-up” section in the Methods for details).

The data that yielded the largest difference (i.e., a performance increase from 0.589 to 0.801, 26%) are visualized in Figure 10. The samples are plotted against the value of their feature (*X*, on the x-axis) and the value of their confound (*C*, on the y-axis), both before subsampling (upper scatter plot) and after subsampling (lower scatter plot). From visual inspection, it appears that the samples rejected by the subsampling procedure (i.e., the samples with the red border) tend to lie relatively close to (or on the “wrong” side of) the decision boundary (i.e., the dashed black line). In other words, subsampling tends to reject samples that are harder to classify based on the data (here, the single feature of *X*). The density plots in Figure 10 show the same effect in a different way: while the difference in the mode of the distribution of the confound (*C*) is reduced after subsampling (i.e., the density plots parallel to the y-axis), the difference in the mode of the distribution of the data (*X*) is actually increased after subsampling (i.e., the density plots parallel to the x-axis).

**Figure 10.**
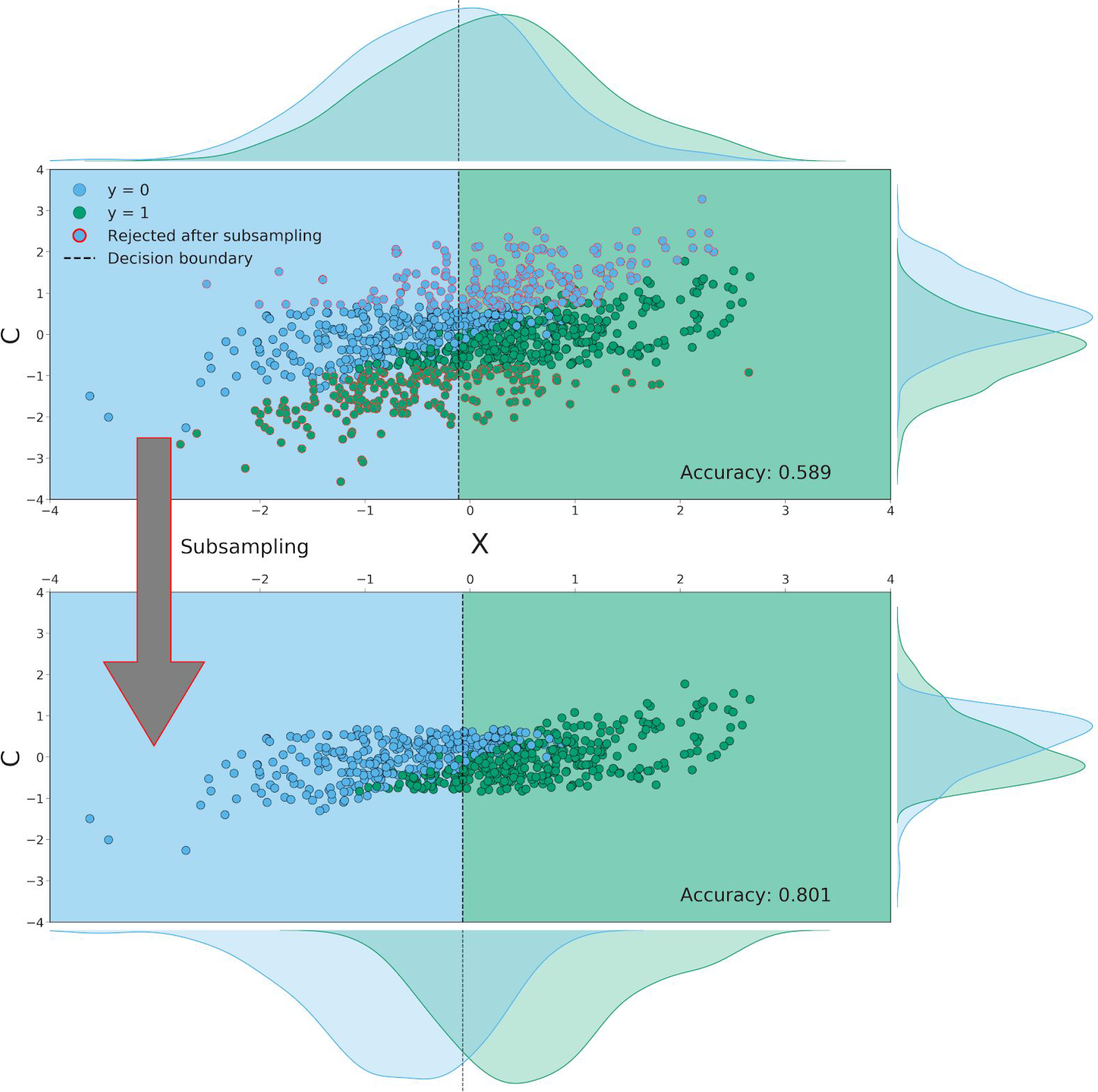
Both scatterplots visualize the relationship between the data (*X* with *K* = 1, on the x-axis), the confound (*C*, on the y-axis) and the target (*y*). Dots with a red border in the upper scatterplot are samples that are rejected in the subsampling process; the lower scatterplot visualizes the data without these rejected samples. The dashed black lines in the scatterplot represent the decision boundary of the SVM classifier; the color of the background shows how samples in that area are classified (a blue background means a prediction of *y* = 0 and a green background means a prediction of *y* = 1). The density plots parallel to the y-axis depict the distribution of the confound (*C*) for the samples in which *y* = 0 (blue) and in which *y* = 1 (green). The density plots parallel to x-axis depict the distribution of the data (*X*) for the samples in which *y* = 0 (blue) and in which *y* = 1 (green), from which it is clear that subsampling preferentially removes samples close to the decision boundary.

We quantified this effect due to subsampling by comparing the signed distance from the decision boundary (i.e., the dashed line in the upper scatter plot) between the retained samples and the rejected (subsampled) samples, in which a larger distance from the decision boundary reflects a higher confidence of the classifier’s prediction (see Figure 3 for a visualization of this method). Indeed, we find that samples that are removed by subsampling lie significantly closer to (or on the “wrong” side of) the decision boundary (*M* = − .358, *SD* = .619) than samples that are retained after subsampling (*M* = .506, *SD* = .580), as indicated by a independent samples *t*-test, *t*(998) = 22.32, *p* < 0.001. Also (which follows from the previous observation), samples that are been removed by subsampling are more often classified incorrectly (75% incorrect) than the samples that would have been retained by subsampling (20% incorrect), as indicated by a chi-squared test, χ^2^ = 270.29, *p*< 0.001.

To show that the same effect (i.e., removing samples that tend to be hard to classify) occurs in the empirical data after counterbalancing, we apply the same analysis of comparing the model performance and distance-to-boundary between the retained and rejected samples to the empirical data. Indeed, across all different amounts of voxels (*K*), the retained samples were classified correctly significantly more often (Figure 11A) and had a significantly larger distance to the classification boundary (Figure 11B) than the rejected samples. This demonstrates that the same effect of counterbalancing as shown in the simulated data (i.e., the removal of hard-to-classify samples) likely underlies the increased model performance of the counterbalanced data relative to the baseline model in the empirical data.

In summary, the removal of a subset of observations to correct for the influence of a confound can induce significant bias by removing samples that are harder to classify using the available data (*X*). The bias itself can be subtle (e.g., in our empirical results, the predictive performance falls in a realistic range of predictive performances), and could remain undetected when present. Therefore, we believe that counterbalancing through subsampling the data is an inappropriate method to control for confounds.

**Figure 11.**
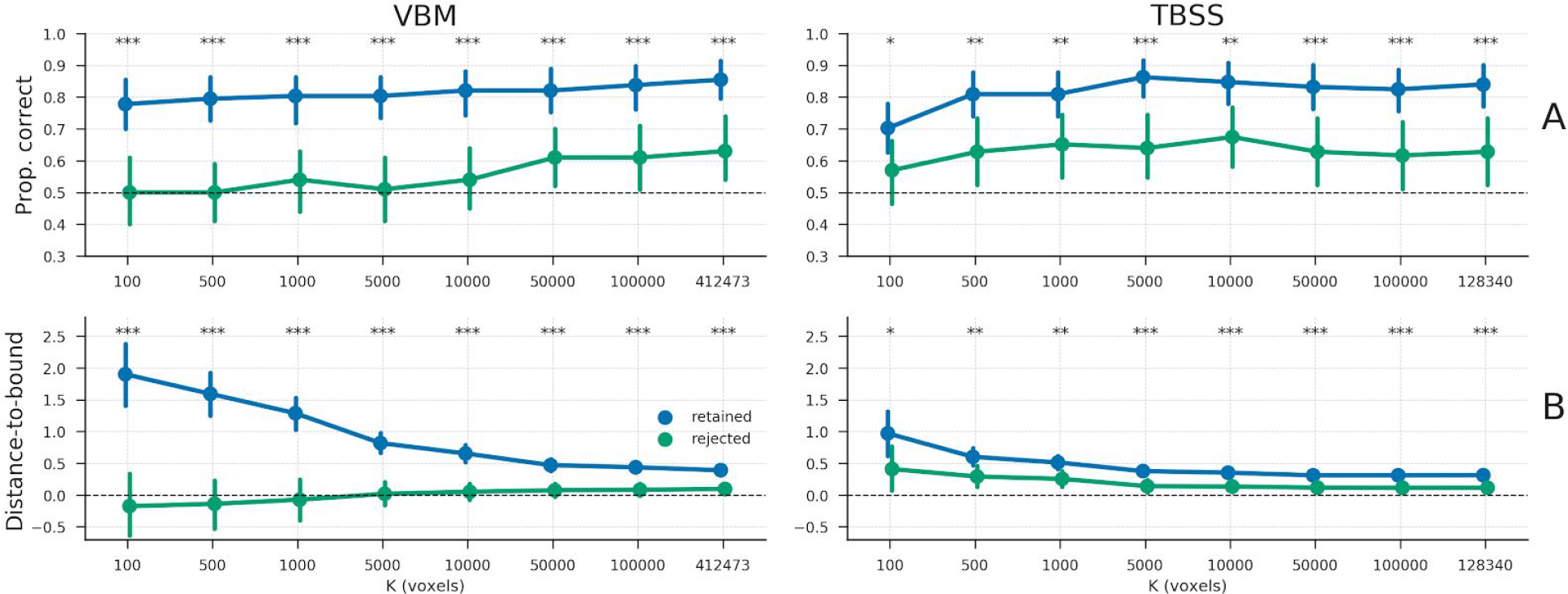
A) The proportion of samples classified correctly, separately for the “retained” samples (blue line) and “rejected” samples (green line); the dashed line represents chance level (0.5). B) The average distance to the classification boundary for the retained and rejected samples; the dashed line represents the decision boundary, with values below the line representing samples on the “wrong” side of the boundary (and vice versa). * = *p* < 0.05, ** = *p* < 0.01, *** *p* < 0.001.

### Whole-dataset confound regression (WDCR)

#### Empirical results

In addition to counterbalancing, we evaluated the efficacy of ‘whole-dataset confound regression’, i.e. regressing out the confound from each feature separately using all samples from the dataset, to control for the of confounds. Compared to the baseline-model, WDCR yielded a strong decrease in performance, even dropping (significantly) below chance for all TBSS-analyses and a subset of the VBM-analyses (see Figure 12).

**Figure 12.**
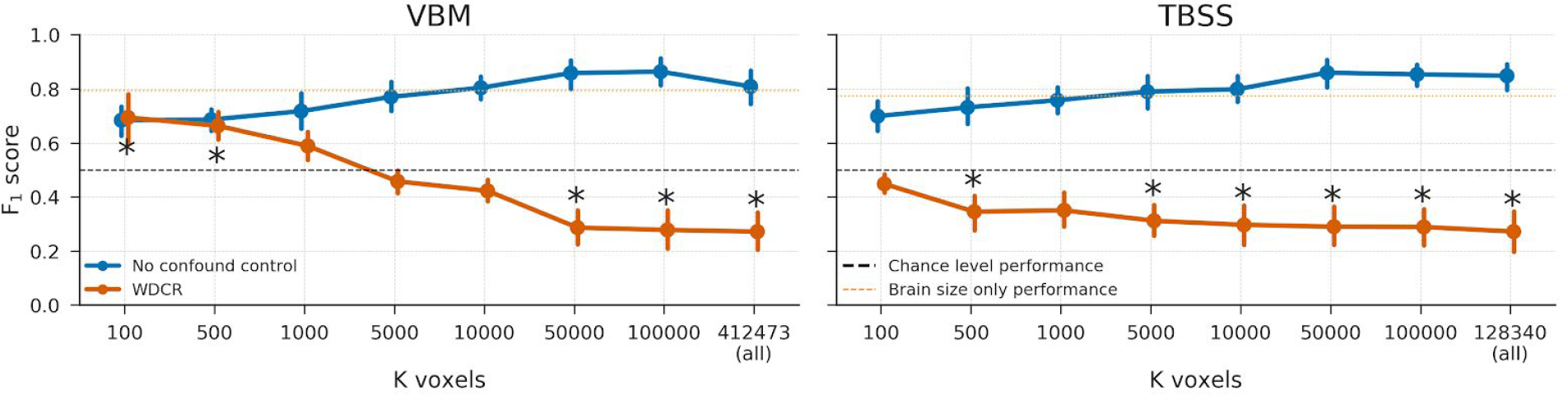
Model performance after WDCR (orange) versus the baseline performance (blue) for both the VBM (left) and TBSS (right) data. Performance reflects the average F_1_ score across 10 folds; error bars reflect 95% confidence intervals across 1000 bootstraps. The dashed black line reflect theoretical chance-level performance (0.5) and the dashed orange line reflects the average model performance when only brain size is used as a predictor; * indicates significant performance (p & 0.001) above/below chance.

This strong (and implausible) reduction in model performance after WDCR is investigated in more detail in the next two sections on the results from the simulations.

#### Generic simulation

In fact, the results from the generic simulation (see Figure 8) show that WDCR accurately corrects for the confound in both the case of data without signal (i.e., when signal *R^2^* = 0.004) and in case of some signal (i.e., when signal *R^2^* = 0.1), as evident from the fact that the performance after WDCR is similar to the reference performance. This result (i.e., plausible performance after confound control) stands in contrast to the results from the empirical analyses, which is why we ran a follow-up simulation to investigate this specific issue, as is reported in the next section.

#### WDCR follow-up simulation

Inspired by the work of Jamalabadi et al. (2016) on below-chance accuracy in decoding analyses, we ran several follow-up analyses to get insight into why WDCR leads to below-chance model performance. As Jamalabadi et al. (2016) show, below-chance model performance occurs when the data contains little signal, which they operationalize using Cohen’s δ (which is proportional to *r_yX_*). In our first follow-up simulation, we sought to refine the explanation of the cause of below-chance model performance by linking it to the observed standard deviation of the empirical distribution of correlations between the data (*X*) and the target (*y*). To do so, we simulate random data (*X*) and a binary target (*y* ∈ {0, 1}) and estimated (per fold) the cross-validated classification accuracy using the standard pipeline described in the methods section. We repeated this process 500 times, yielding 500 data sets. The expected *average* predictive accuracy for each dataset is 0.5, but this varies randomly across folds and iterations. We hypothesized that this variance can be explained by the standard deviation (“width”) of the initial feature-target correlation distribution, *sd*(*r_yX_*) : narrower distributions may yield relatively lower cross-validated classification accuracy than relatively wider feature-target correlation distributions. Indeed, we find that the initial standard deviation of this distribution is significantly correlated with the cross-validated accuracy, *r*(499) = 0.73, *p* < 0.001 (Figure 13A). Importantly, we find that this relationship holds for different values of *N* (see Supplementary Figure 1), for different sizes of the test-set (see Supplementary Figure 2), and for different sizes of *K* (see Supplementary Figure 3).

**Figure 13.**
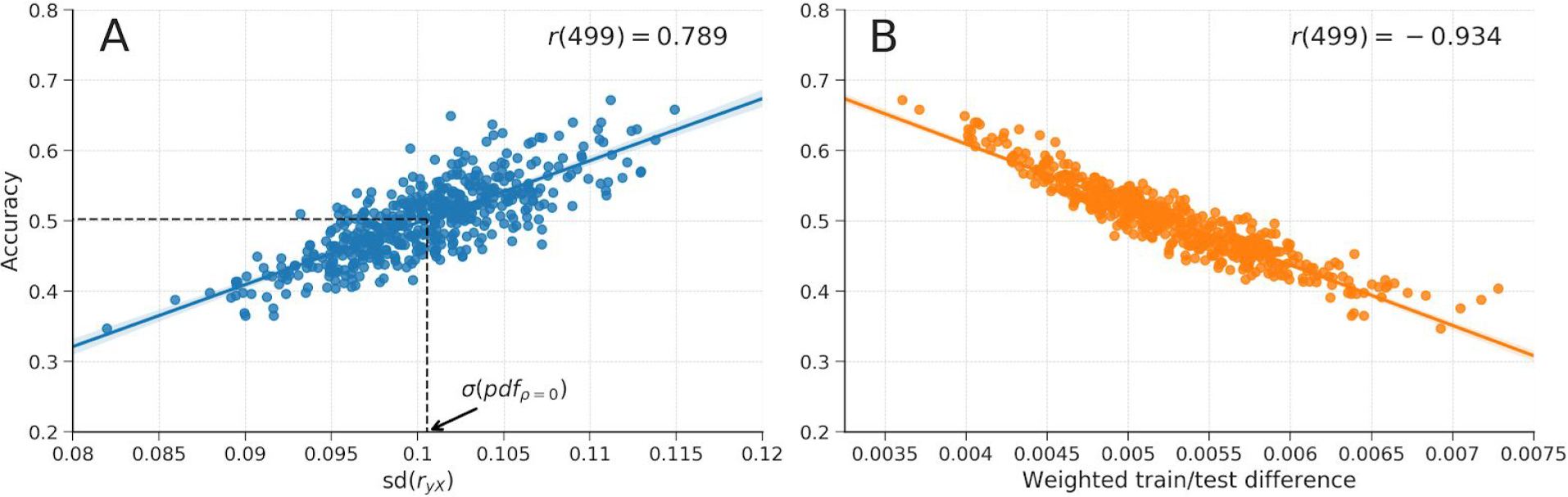
A) The relationship between the standard deviation of the distribution of feature-target correlations, *sd*(*r_yX_*), and accuracy across iterations of cross-validated classification analyses of null-data. The vertical dashed line represents the standard deviation from the sampling distribution parameterized with ρ = 0 and *N* = 100 (i.e., the same parameters used to generate the null-data); the horizontal dashed line represents the expected accuracy for data with this standard deviation based on the regression line estimated from the data across simulations (see Supplementary Figure 1 for the same plot with different values for *N*). B) The relationship between the weighted difference between feature-target correlations in the train and test set and accuracy.

This observation, then, begs the question: *why* do narrower-than-chance correlation distributions lead to below-chance accuray? In follow-up analyses of the data in Figure 13A, we found that relatively narrow distributions of feature-target correlations induce what is known in the machine learning literature as “dataset shift”, which describes the phenomenon of a change in feature-target relationship between the train and test set (Jamalabadi et al., 2016; Quionero-Candela, Sugiyama, Schwaighofer, & Lawrence, 2009). An example of “dataset shift” is observing a positive correlation of 0.1 between a particular feature and the target in the train set while observing a negative correlation of −0.1 for this same feature is the test set. We quantified the effect induced by “dataset shift” as the average (across features *j* = 1, …, *K*) difference between the train-set feature-target correlation (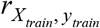) *train train*and test-set feature-target correlation (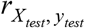), weighted by coefficients^13^ assigned to the *test test* feature by the classifier (*w*). Formally, we estimate dataset shift (*d*^̂^*s*) as follows:

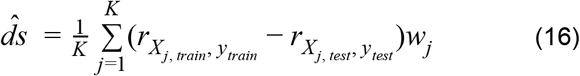

Indeed, the correlation between this particular operationalization of “dataset shift” and accuracy across simulations is highly significant, *r*(499) = −0.934 (Figure 13B).

Having established the relation between the standard deviation of the initial feature-target correlation distribution and (below-chance) accuracy, we followed up our simulation by investigating specifically the effect of WDCR on the standard deviation of the correlation distribution. We investigated this by simulating data with different strengths of the correlation between the confound and the target (*r_Cy_*) and the number of features (*K*). In Figure 14A, it is clear that, while the expected chance level is 0.5 in all cases, model performance quickly drops below chance for increasing correlations between the target and the confound and for increasing numbers of features, and even leading to a model performance of 0% correct when the the confound is perfectly correlated with the target and when using 1000 features. Figure 14C shows that, indeed, higher *r_Cy_* values lead to smaller widths of the correlation distribution, which is shown in Figure 14D to yields relatively lower accuracy scores.

**Figure 14.**
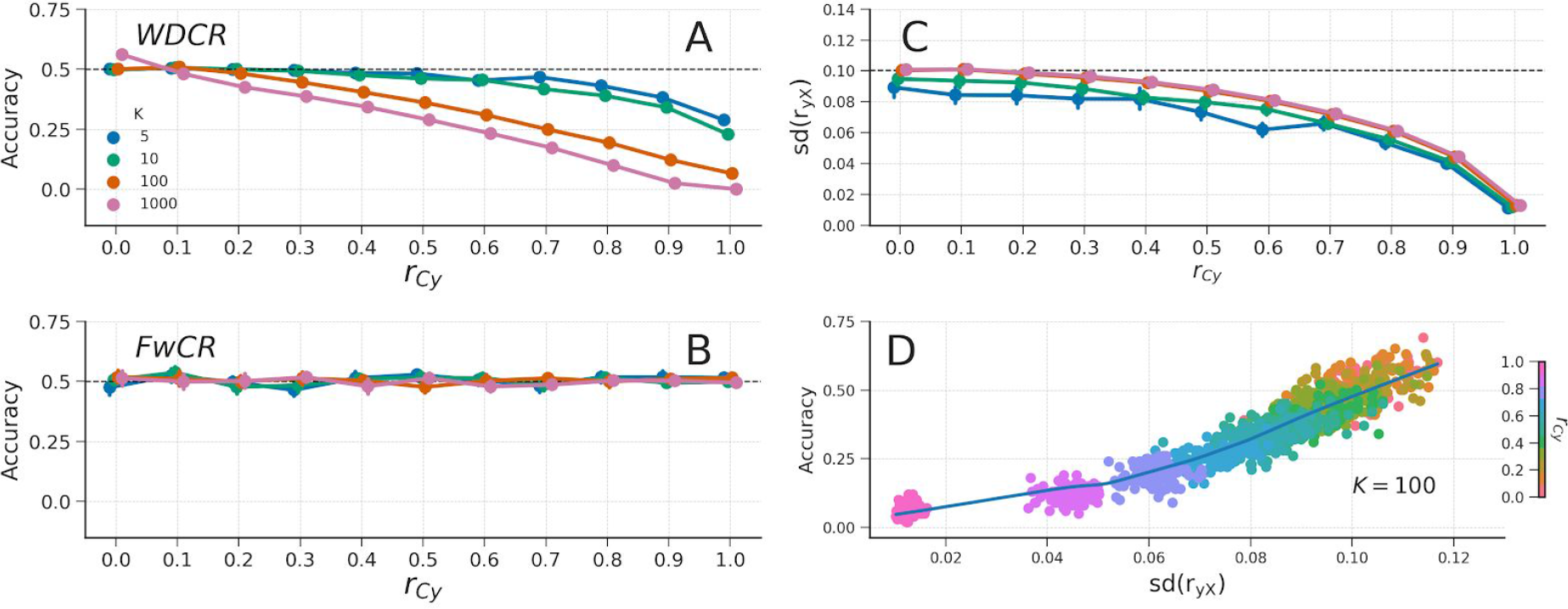
A) The effect of WDCR on data varying in the correlation of the confound with the target (*r_Cy_*; x-axis) and the number of features (*K*; different lines). B) The effect of FwCR on data varying in the correlation of the confound with the target and the number of features. The dashed black line represents chance model performance in subplots A and B. C) The relation between the correlation of the confound with the target (*r_Cy_*) and the standard deviation of the feature-target correlation distribution, *sd*(*r_yX_*) for the WDCR data. The dashed black line represents the standard deviation of the correlation distribution predicted by the sampling distribution. D) The relation of the standard deviation of the correlation distribution and accuracy for the WDCR data (only shown for the data when *K* = 100; see Supplementary Figure 4 for visualizations of this effect for different values of *K*). The data depicted in all panels are null-data (see WDCR/FwCR follow-up simulation for simulation details).

In summary, our simulations show that below-chance accuracy is accurately predicted by the standard deviation (i.e., “width”) of the distribution of empirical feature-target correlations and that WDCR reduces this standard deviation, which explain why the empirical analyses yielded below-chance model performance (especially for larger amounts of voxels).

### Foldwise confound regression (FwCR)

#### Empirical results

As the results from the empirical analyses and simulations suggest, WDCR appear problematic because of the partitioning of the dataset into a separate train-set and test-set after confound regression. As such, our proposed foldwise confound regression (FwCR) suggest to move the confound regression procedure inside the cross-validation loop and thus also cross-validating this step. As expected, compared to the baseline model (i.e., no confound control), the results from the empirical analyses using FwCR show reduced (but not below-chance) model performance for both VBM and TBSS and all different number of voxels (see Figure 15). All performance estimates (for all amounts of voxels and in both VBM and TBSS) are significant (*p* < 0.001).

**Figure 15.**
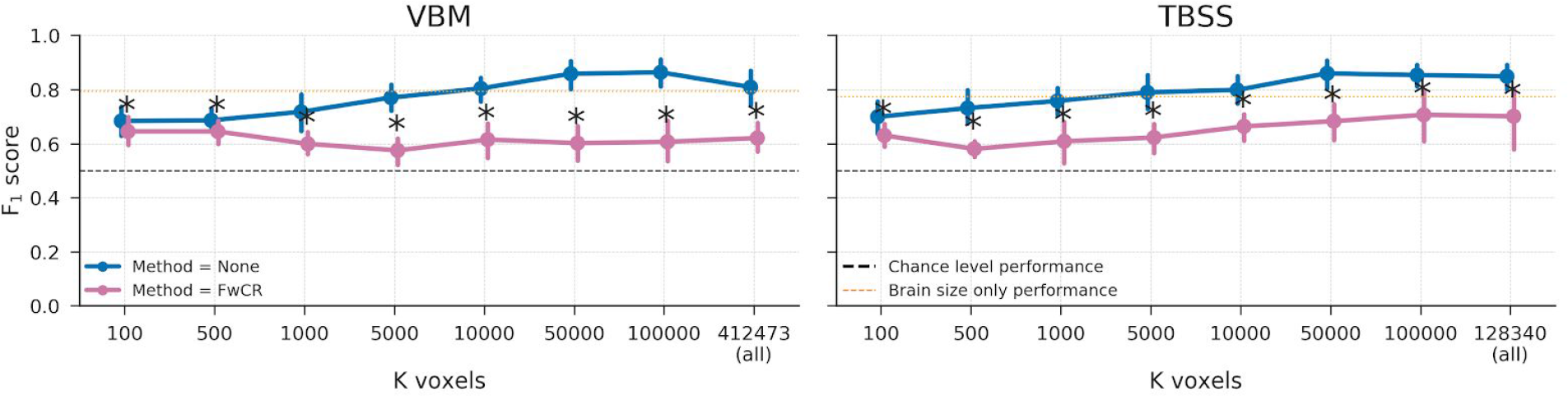
Model performance after FwCR (pink) versus the baseline performance (blue) for both the VBM (left) and TBSS (right) data. Performance reflects the average F_1_ score across 10 folds; error bars reflect 95% confidence intervals across 1000 bootstraps. The dashed black line reflect theoretical chance-level performance (0.5) and the dashed orange line reflects the average model performance when only brain size is used as a predictor. * indicates significant performance (p < 0.001) above chance.

#### Generic simulation

Similar to WDCR, FwCR yields plausible and unbiased model performance (see Figure 8, pink line).

#### FwCR follow-up simulation

When applied to the simulated null-data as described in the “WDCR follow-up simulation” section, FwCR yields model performance scores at chance level across all levels of the confound-target correlation and different amount of features (see Figure 14B).

#### Summary methods for confound control

In this section, we have shown the effects of different method to control confounds (counterbalancing, WDCR, and FwCR) on empirical MRI data and simulated data (see Figure 16). Counterbalancing is clearly unable to correctly control for confounding influences, which is putatively caused by indirect circularity in the analysis process due to subsampling. Confound regression shows an expected drop in model performance (but not below chance-level), given that the confound regression step is properly cross-validated (i.e., the FwCR version).

**Figure 16.**
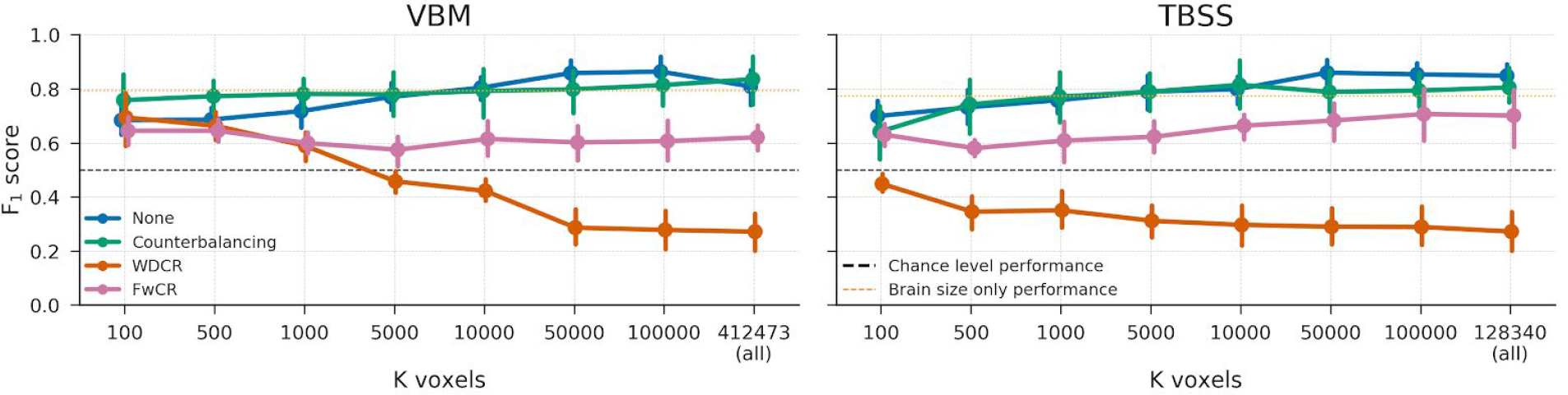
An overview of the empirical results on the four different confound methods: None, Counterbalancing, WDCR, and FwCR.

## DISCUSSION

Decoding has become a popular alternative to univariate analyses of neuroimaging data. This analysis approach, however, inherently suffers from ambiguity in the source of information picked up by the decoder (Naselaris & Kay, 2015). Given that one is interested in model interpretability rather than merely accurate prediction (Hebart & Baker, 2017), one should strive to control for alternative sources of information (i.e., other than the target-of-interest) that might drive decoding. Effectively controlling for these alternative sources of information, or *confounds*, helps in disambiguating decoding models. In this article, we reviewed and tested two generic, broadly applicable methods that aim to control for confounds in decoding analyses: (post-hoc) counterbalancing and confound regression. Additionally, we proposed a third method (a modification of traditional confound regression) that, unlike the other two methods, effectively and in an unbiased way controls for confounds.

In both the empirical data and simulations, we found that neither (post-hoc) counterbalancing nor (whole-dataset) confound regression yielded plausible and unbiased model performance estimates. First, we found that post-hoc counterbalancing leads to *optimistic* (i.e., positively biased) model performance estimates, which is caused by removing samples that are hard to classify during the subsampling process. Because this subsampling process is applied to the entire dataset at once (i.e., it is not cross-validated), it can be seen as a form of indirect circular analysis (Kriegeskorte et al., 2009), in which the data itself is used to inform analysis decisions, and is thus bound to yield optimistic generalization estimates. Second, our initial evaluation of confound regression, which was applied on the entire dataset (“WDCR”), yielded *pessimistic* (i.e., negatively biased) and even significantly below-chance model performance estimates. Extending previous research (Jamalabadi et al., 2016), we show that this negative bias occurs when the “signal” in the data (operationalized as the width of the feature-target correlation distribution) is lower than would be expected by chance, which we link to the sampling distribution of the Pearson correlation coefficient. Importantly, we show that WDCR systematically narrows the width of the correlation distribution — and thus leads to lower model performance — which is exacerbated by higher correlations by the target and the confound and more features.

To overcome the negative bias observed in WDCR, we propose to cross-validate the confound regression procedure (which we call “Foldwise Confound Regression”, FwCR). We show that this method yields plausible model performance in the empirical analyses (i.e., significantly above-chance model performance) and nearly unbiased model performance in the simulations for different datasets varying in the amount of features (*K*) and the strength of the confound (*r_Cy_*). As FwCR can be used with any type of data (electrophysiology, MRI, and even behavioral data), it represents a universal method to disambiguate decoding analyses^14^.

### Relevance and consequences for previous and future research

We believe our results have implications for not only *post-hoc* counterbalancing, but counterbalancing in general. In both behavioral research (Wacholder, Silverman, McLaughlin, & Mandel, 1992) and neuroimaging research (Görgen et al., 2017), a priori counterbalancing (or case-control ‘matching’) is common to avoid effects of confounds. However, as we show in the current study, this may unintentionally remove samples that are harder to predict, especially when there is little shared variance between the confound and the other predictors (i.e., when there is low confound *R^2^*). Because, conceptually, this represents a form of circular analysis, counterbalancing — regardless of whether it is applied a priori or post-hoc — is bound to yield overly optimistic model performance estimates. Another way of interpreting this finding, more in terms of classical frequentist statistics, is that the model performance estimates are based on a sample that is not representative of the population (see also Sedgwick, 2013). As a result, out-of-sample predictive performance drops significantly, in our case even to chance level.

Since counterbalancing does not show any positive bias in model performance when there is no signal at all (i.e., signal *R*^2^ = 0), one could argue that any observed significant above-chance effect, while positively biased in terms of effect magnitude, can be interpreted as reflecting the fact that there must be signal in the data in the first place. However, we argue against this interpretation for two reasons. First, any above-chance predictive performance of models fitted after subsampling is not only positively biased, but also does not cross-validate to the rejected samples. That is, the model picks up relations between features and target that are only present in the subsample, and not in the samples left out of the analysis. As a result, it is questionable whether (and if so, how) the model should be interpreted—after all, the rejected samples were validly drawn from the population of interest. Second, any possible *absence* of above-chance model performance after subsampling can neither be interpreted as evidence for an absence of a true effect, since the subsampling procedure necessarily leads to a (often substantial) power loss. It could still well be that in the original sample there was a true relation between features and target. Thus, interpretation of modelling efforts after subsampling is troubled in case of both presence and absence of above-chance model performances.

In contrast to post-hoc counterbalancing, confound regression in its uncross-validated form (i.e., WDCR) has been applied widely in the context of decoding analyses (Dubois et al., 2017; Kostro et al., 2014; Rao et al., 2017; Todd et al., 2013). Indeed, the first study that systematically investigated the effect of confounds in decoding analyses (Todd et al., 2013) used WDCR to account for the confounding effect of reaction times (RT) on decoding of rule representations and found that WDCR completely eliminated the effect found when not controlling for RT. This observation, however, can potentially be explained by the negative bias induced by WDCR. This possible explanation is corroborated by a follow-up study that similarly looked into RT confounding decoding of rule representations (Woolgar et al., 2014), who did not use WDCR but accounted for RT confounding by including it as a covariate during the pattern estimation procedure (see “Control for confounds during pattern estimation” section), which in contrast to the study by Todd et al. yielded widespread significant decoding. Moreover, while not specifically investigated here, we expect a similar negative bias to occur when a confound is removed from a continuous target variable using WDCR - which may offer an explanation for the null-finding of (Dubois et al., 2017), who fail to decode personality characteristics from resting-state fMRI.

Interestingly, a technique related to confound regression is recently gaining popularity in studies using representational similarity analysis (RSA) to analyze neural data (Kriegeskorte, Mur, & Bandettini, 2008). In the context of RSA, the explained variance in the neural data is often partitioned into different (model-based) feature sets (i.e., sources of information), which allows one to draw conclusions about the *unique* influence of each source of information (see, e.g., Groen et al., 2018; Hebart, Bankson, Harel, Baker, & Cichy, 2018; Ramakrishnan, Scholte, Groen, Smeulders, & Ghebreab, 2014). Specifically, variance partitioning in RSA is done by removing the variance from the representational dissimilarity matrix (RDM) based on the feature set that needs to be controlled for. Notably, the variance of the RDMs that are not of interest can be removed from only the neural RDM (Hebart et al., 2018; Ramakrishnan et al., 2014) or both from the neural RDM *and* the RDM-of-interest (Groen et al., 2018). While the context is different, the underlying technique is identical to confound regression as described and evaluated in this article. Importantly, the studies employing this variance partitioning technique (Groen et al., 2018; Hebart et al., 2018; Ramakrishnan et al., 2014) similarly report plausible model performances after confound regression (i.e., relatively lower but not below-chance performance), corroborating our results with (foldwise) confound regression. Note that the distinction between WDCR and FwCR in the context of most RSA studies (including the aforementioned studies) is irrelevant, as representational similarity analyses are not commonly cross-validated. However, recently, some have proposed to use cross-validated distance measures (such as the cross-validated Mahalanobis distance; Guggenmos, Sterzer, & Cichy, 2018; Walther et al., 2016) in representational similarity analyses, which could suffer from negative bias when combined with (not cross-validated) variance partitioning similar to what we observed with WDCR in the context of decoding analyses.

The importance of proper confound control is highlighted by the empirical question we address. Without any prediction pipeline optimization, we were able to predict gender with a model performance up to approximately 0.85 without confound control. This is in line with reports from various other studies (Del Giudice et al., 2016; Rosenblatt, 2016; Sepehrband et al., 2018). However, this predictive performance is driven by a mixture two sources of information: global and local differences in brain structure. With confound control, however, we show that predictive performance using only local differences lies around 70%—a substantial drop in performance. Especially because the remaining predictive performance is lower than predictive performance using only brain size, we argue that proper confound control may lead one to draw significantly different conclusions about the differences in brain structure between men and women. For the debate on sexual dimorphism, it is thus extremely important to take global brain size into account in the context of decoding analyses (as has been similarly recommended for mass-univariate analyses; Barnes et al., 2010).

### Practical recommendations

As indicated by the title of this article, we will now outline some practical recommendations for dealing with confounds in decoding analyses of neuroimaging data. First, one can only control for confounding variables when it these variables are actually *known* to confound the decoding analysis (i.e., they are correlated to the target and encoded in the neuroimaging data). To detect confounds, we recommend using the “same analysis approach” outlined by Görgen and colleagues (2017). In short, this method recommends trying to predict the target variable using your confound(s) as predictive features (like we did when using only brain size to predict gender); in case of significant above-chance decoding performance, and assuming the confounds are actually encoded in the neuroimaging data, the hypothesized confounds will most likely influence the actual decoding analysis. While in the current article we focused on simple univariate confounding effects (i.e., confounding by a single variable), the same analysis approach is not limited to detecting univariate confounds — it facilitates detecting multivariate (i.e., confounding by multiple variables) or interaction effects (i.e., confounding by interaction effects between variables) as well. For example, if one hypothesizes that the target variable is related to the interaction between confound *C*_1_ and *C*_2_ (i.e., *C*_1_ × *C*_2_), one can simply use the interaction term as the potential confound in the same analysis approach to evaluate the potential confounding influence.

Once the specific confound terms has been identified, we recommend regressing out the confound from the data in a cross-validated manner (i.e., FwCR). Specifically, we recommend including confound regression as the first step in your decoding pipeline to avoid the effect of confounds on other operations in the pipeline (such as univariate feature selection; Chu et al., 2012). In this article, we used ordinary least squares (OLS) regression to remove the influence of confounds from the data, for reasons of parsimony (we had no reason to assume a non-linear relation between the confound and the neuroimaging data, so we chose the simplest model) and computational efficiency (OLS can be easily vectorized across all features (*j* = 1, …, *K*), making it a fast confound regression model). However, FwCR is not limited in any way to simple models; any model of arbitrary complexity can be used. We argue that the principle of parsimony should be guiding when choosing a confound regression model: simple models should be preferred, unless one has clear reasons to suspect that a simple model is not adequate. In such cases a different, nonlinear model can be chosen (see e.g., Abdulkadir et al., 2014; Kostro et al., 2014).

## CONCLUSIONS

In general, we believe that the contributions of the current study are twofold. First and foremost, it provides a systematic evaluation of two widely applicable methods to control for confounds and shows that only a single method (“foldwise confound regression”) yields plausible and unbiased results. The results from this evaluation hopefully prevents researchers from using counterbalancing and (whole-dataset) confound regression, which we show may introduce (unintended) biases. Moreover, we open-sourced all analyses and preprocessed data (https://github.com/lukassnoek/MVCA) and provide a simple implementation for foldwise confound regression that interfaces with the popular scikit-learn package in Python. Second, we believe that this study improves understanding of the elusive phenomenon of below-chance accuracy (building on previous work by Jamalabadi et al., 2016). In general, we hope that this study helps researchers in gaining more insight into their decoding analyses by providing a method that disentangles the contributions of different sources of information that may by encoded in their data.

## CHANGELOG

### Version 23 April, 2018

Below, the changes with respect to the previous version on Biorvix (from March 28, 2018) are listed.

- Add a paragraph about concept-response curve (Alizadeh et al., 2017) as a method to deal with confounds (in “Include confounds in model” section);
- Change sign in Figure 2 (was >, but should be <);
- Remove references to work of the Gallant lab upon request;
- Add paragraph in discussion about why you shouldn’t use counterbalancing, even when there is no signal;
- Add new paragraph in discussion about practical recommendations with respect to confound control (including information about the scope of confound regression)

## Supplementary Materials

**This document includes**

Supplementary Figures 1-4

**Supplementary Figure 1.**
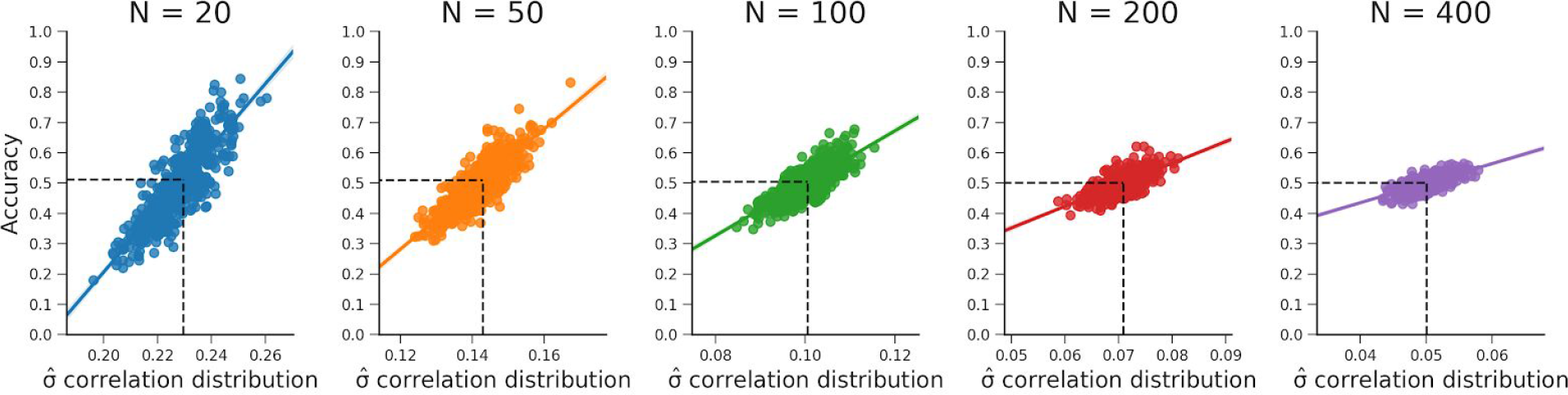
These plots show that the relationship between the standard deviation of the empirical feature-target correlation distribution, *sd*(*r_yX_*), and accuracy holds for different samples sizes (i.e., values for *N*). Note that the predicted accuracy based on the standard deviation expected from the sampling distribution is at 0.5 for every plot. The data were generated in the same manner as reported in the “WDCR follow-up” section.

**Supplementary Figure 2.**
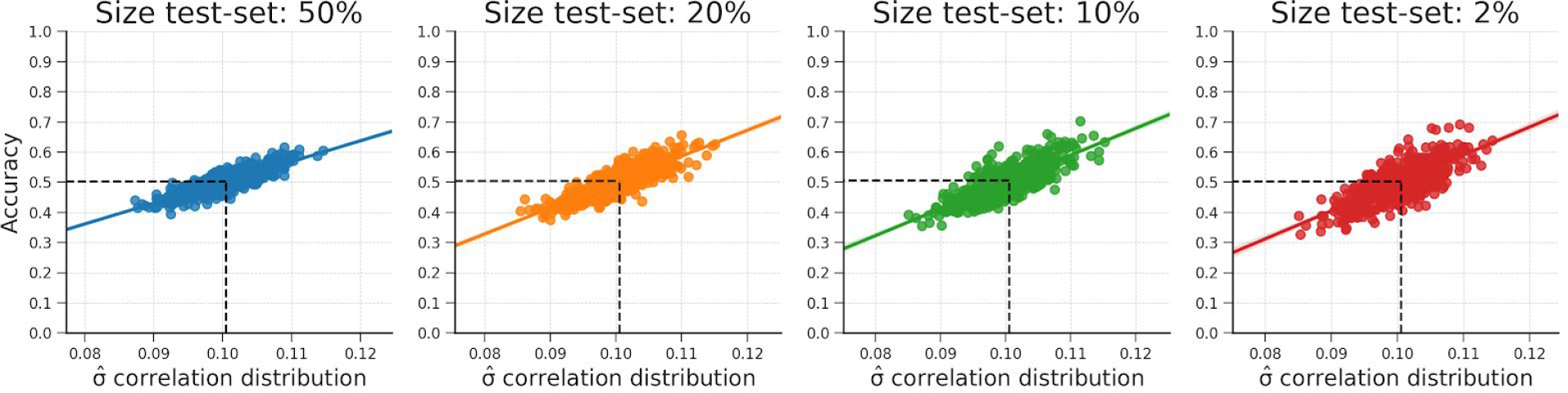
These plots show that the relationship between the standard deviation of the empirical feature-target correlation distribution, *sd*(*r_yX_*), and accuracy also holds for sizes of the test-set (replicating results from Jamalabadi et al., 2016). Note that the predicted accuracy based on *sd*(*r_yX_*) is again at 0.5 for every plot. The data were generated in the same manner as reported in the “WDCR follow-up” section.

**Supplementary Figure 3.**
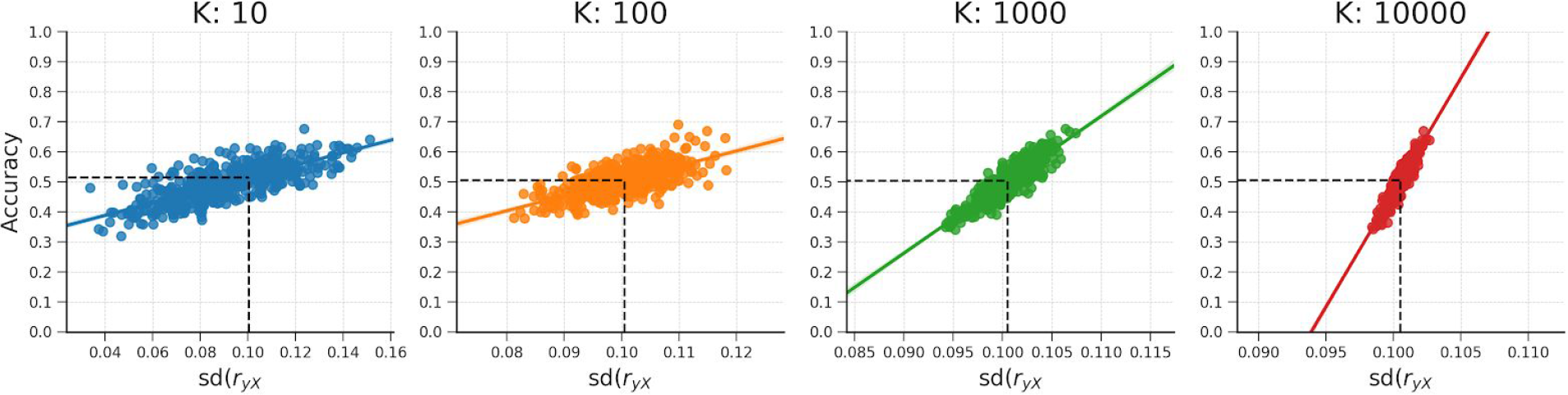
These plots show that the relationship between the standard deviation of the empirical feature-target correlation distribution, *sd*(*r_yX_*), and accuracy also holds for different amounts of features (*K*). Note that the predicted accuracy based on *sd*(*r_yX_*) is approximately at 0.5 for every plot. The data were generated in the same manner as reported in the “WDCR follow-up” section.

**Supplementary Figure 4.**
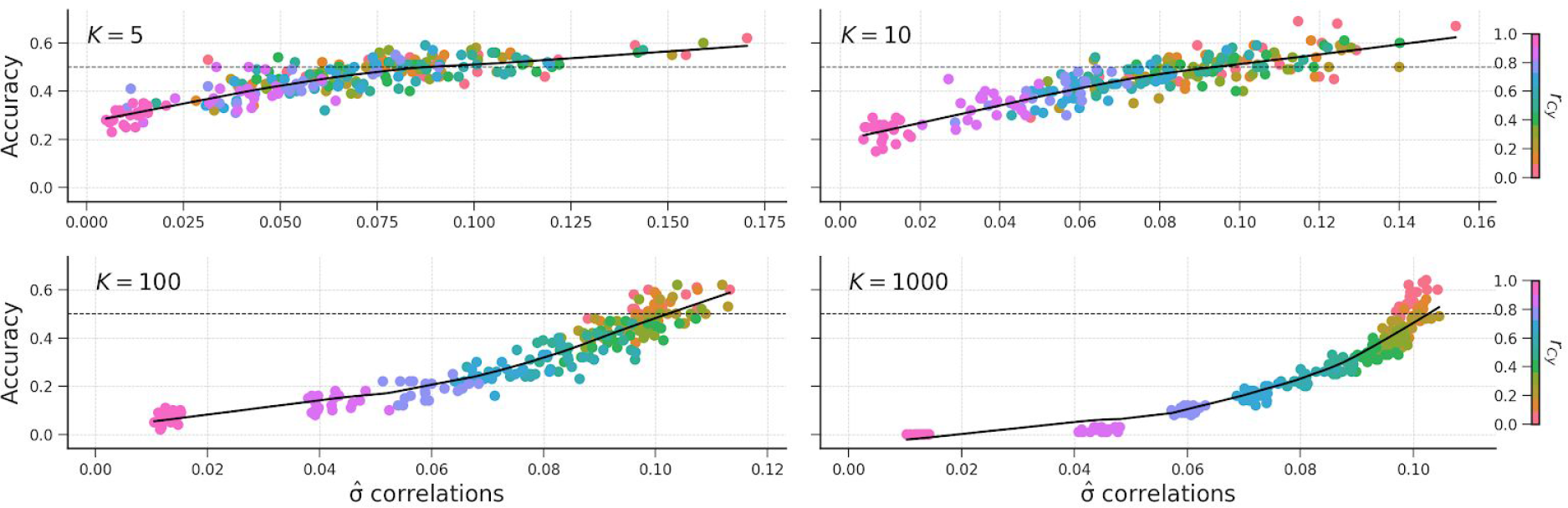
The relation of the standard deviation of the correlation distribution and accuracy for different values of *K*.

## Footnotes

1 However, if accurate prediction is the *only* goal in this scenario, we would argue that there are probably easier and less expensive methods than neuroimaging to predict a participant’s gender.

2 Note that our definition of a confound assumes that the primary aim of a decoding analysis in *interpretability* rather than mere *prediction*. In fact, when one is interested in accurate prediction only, any source of information covarying with the target (given that this effect is also present in the population) is actually beneficial for model performance and is thus often *not* regarded as problematic (see for a different definition, e.g., Rao et al., 2017, p. 40).

3 In the context of behavioral data, a priori counterbalancing is often called ‘matching’ or a employing a ‘case-control design’ (Cook, Campbell, & Shadish, 2002).

4 Note that the counterbalancing process is the same for both univariate (activation-based) studies and decoding studies, but the direction of analysis is reversed in univariate (e.g., gender → brain) and decoding studies (e.g., brain → gender). As such, in univariate studies the confound (e.g., brain size) is counterbalanced with respect to the *predictor*(*s*) (e.g., gender) while in decoding studies the confound (e.g., brain size) is counterbalanced with respect to the *target* (e.g., gender).

5 Parameter estimates only reflect unique variance when ordinary least squares is used to find the model parameters. Other (regularized) linear models, such as ridge regression or LASSO, are not guaranteed to yield parameters with unique variance.

6 Note that *X* and *C*, here, refer to (usually HRF-convolved) predictors of the time series signal (*s*) for a *single* voxel. In the rest of the article, *X* and *C* refer to features that are defined across samples (not time).

7 Note that, technically, one could use the “Control for confounds during pattern estimation” method in electrophysiology, by first fitting a univariate model explaining the neuroimaging data (*X_j_* for j = 1, …, *K*) as a function of *both* the target (*y*) and the confound (*C*) and subsequently only using the parameter estimates of the target-predictor (*β_y_*) as patterns in the subsequent decoding analysis; this is, in fact, equivalent to the “confound regression” technique, discussed in the next section.

8 While uncommon in decoding analyses, confound regression is more common as a tool for variance partitioning in neuroimaging studies using representational similarity analysis (Groen et al., 2018; Hebart, Bankson, Harel, Baker, & Cichy, 2018), which is discussed in more detail in the discussion.

9 Note that we did not discuss studies that implement a different confound regression procedure (e.g., Abdulkadir, Ronneberger, Tabrizi, & Klöppel, 2014; Dukart, Schroeter, Mueller, et al., 2011), in which confound regression is only estimated on the samples from a single class of the target variable (e.g., in our gender decoding example, this would mean that confound regression models are only estimated on the data from male, or female, subjects). As this form of confound regression does not disambiguate the sources of information driving the decoder, it is thus not discussed further in this article.

10 For continuous confounds, it is practically impossible achieve a correlation with the target of *exactly* zero, which is the reason we subsample until it is smaller than a prespecified threshold. For categorical confounds, however, a correlation between the confound and the target is possible (this amounts to equal proportions of levels of *c* within each class of *y*; (Görgen, Hebart, Allefeld, & Haynes, 2017), even *necessary*, because it is impossible to find a (K-fold) cross-validation partitioning in which each split is counterbalanced w.r.t. the confound if the correlation *in the entire dataset* between the target and the confound is not zero.

11 Note that this is more strict than the conventionally used threshold (α = 0.05), but given that decoding analyses are often more sensitive to signal (whether it is confounded or true signal) in the data, we choose to err on the safe side and counterbalance the data using a relatively strict threshold of α = 0.1.

12 Note that *plausible* null data does not reflect a signal *R*^2^ of 0, because this statistic is biased towards values larger than 0 (because it represents a squared number) when dealing with noisy data, hence our choice of *signal R^2^ = 0.004*.

13 We weigh the train-test difference in feature-target correlations with the classifier weights to capture the importance of each feature in the model.

14 While FwCR offers an unbiased method to control for confounds in decoding analyses, it is limited to confounding effects that are linearly encoded in the neuroimaging data (because, here, we use OLS to regress out the confound from the data). Theoretically, if one would use a non-linear model (such as radial-basis SVMs or decision trees) in a decoding analysis, this model could potentially pick up confounded signal that is non-linearly related to the target variable. However, this issue could be mitigated by using non-linear models to regress out the confound from the data (as done in, e.g., Abdulkadir et al., 2014; Kostro et al., 2014).

## REFERENCES

Abdulkadir, A., Ronneberger, O., Tabrizi, S. J., & Klöppel, S. (2014). Reduction of confounding effects with voxel-wise Gaussian process regression in structural MRI. In 2014 International Workshop on Pattern Recognition in Neuroimaging (pp. 1–4).

Abraham, A., Pedregosa, F., Eickenberg, M., Gervais, P., Mueller, A., Kossaifi, J., … Varoquaux, G. (2014). Machine learning for neuroimaging with scikit-learn. Frontiers in Neuroinformatics, 8, 14.

Allefeld, C., Görgen, K., & Haynes, J.-D. (2016). Valid population inference for information-based imaging: From the second-level t-test to prevalence inference. NeuroImage, 141, 378–392.

Bangalore, S. S., Prasad, K. M. R., Montrose, D. M., Goradia, D. D., Diwadkar, V. A., & Keshavan, M. S. (2008). Cannabis use and brain structural alterations in first episode schizophrenia—a region of interest, voxel based morphometric study. Schizophrenia Research, 99(1), 1–6.

Barnes, J., Ridgway, G. R., Bartlett, J., Henley, S. M. D., Lehmann, M., Hobbs, N., … Fox, N. C. (2010). Head size, age and gender adjustment in MRI studies: a necessary nuisance? NeuroImage, 53(4), 1244–1255.

Bzdok, D. (2017). Classical Statistics and Statistical Learning in Imaging Neuroscience. Frontiers in Neuroscience, 11, 543.

Carlson, T. A., & Wardle, S. G. (2015). Sensible decoding. NeuroImage, 110, 217–218.

Chekroud, A. M., Ward, E. J., Rosenberg, M. D., & Holmes, A. J. (2016). Patterns in the human brain mosaic discriminate males from females. Proceedings of the National Academy of Sciences of the United States of America, 113(14), E1968.

Chu, C., Hsu, A. L., Chou, K. H., Bandettini, P., Lin, C., & Alzheimer’s Disease Neuroimaging Initiative. (2012). Does feature selection improve classification accuracy? Impact of sample size and feature selection on classification using anatomical magnetic resonance images. Neuroimage, 60(1), 59-70.

Cook, T. D., Campbell, D. T., & Shadish, W. (2002). Experimental and quasi-experimental designs for generalized causal inference. Houghton Mifflin Boston.

Craddock, R. C., Holtzheimer, P. E., 3rd, Hu, X. P., & Mayberg, H. S. (2009). Disease state prediction from resting state functional connectivity. Magnetic Resonance in Medicine: Official Journal of the Society of Magnetic Resonance in Medicine / Society of Magnetic Resonance in Medicine, 62(6), 1619–1628.

Cuingnet, R., Gerardin, E., Tessieras, J., Auzias, G., Lehéricy, S., Habert, M.-O., … Alzheimer’s Disease Neuroimaging Initiative. (2011). Automatic classification of patients with Alzheimer’s disease from structural MRI: a comparison of ten methods using the ADNI database. NeuroImage, 56(2), 766–781.

Davis, T., LaRocque, K. F., Mumford, J. A., Norman, K. A., Wagner, A. D., & Poldrack, R. A. (2014). What do differences between multi-voxel and univariate analysis mean? How subject-, voxel-, and trial-level variance impact fMRI analysis. NeuroImage, 97, 271–283.

Del Giudice, M., Lippa, R. A., Puts, D. A., Bailey, D. H., Bailey, J. M., & Schmitt, D. P. (2016). Joel et al.’s method systematically fails to detect large, consistent sex differences. Proceedings of the National Academy of Sciences of the United States of America, 113(14), E1965.

Diedrichsen, J., & Kriegeskorte, N. (2017). Representational models: A common framework for understanding encoding, pattern-component, and representational-similarity analysis. PLoS Computational Biology, 13(4), e1005508.

Dixon, L. (1999). Dual diagnosis of substance abuse in schizophrenia: prevalence and impact on outcomes. Schizophrenia Research, 35 Suppl, S93–S100.

Douaud, G., Smith, S., Jenkinson, M., Behrens, T., Johansen-Berg, H., Vickers, J., … James, A. (2007). Anatomically related grey and white matter abnormalities in adolescent-onset schizophrenia. Brain: A Journal of Neurology, 130(Pt 9), 2375–2386.

Dubois, J., Galdi, P., Han, Y., Paul, L. K., & Adolphs, R. (2017, November 7). Predicting personality traits from resting-state fMRI. *bioRxiv*. https://doi.org/10.1101/215129

Dukart, J., Schroeter, M. L., Mueller, K., Initiative, A. D. N., & Others. (2011). Age correction in dementia–matching to a healthy brain. PloS One, 6(7), e22193.

Gilron, R., Rosenblatt, J., Koyejo, O., Poldrack, R. A., & Mukamel, R. (2017). What’s in a pattern? Examining the type of signal multivariate analysis uncovers at the group level. NeuroImage, 146, 113–120.

Glezerman, M. (2016). Yes, there is a female and a male brain: Morphology versus functionality. Proceedings of the National Academy of Sciences, 113(14), E1971–E1971.

Goldstein, J. M., Seidman, L. J., Horton, N. J., Makris, N., Kennedy, D. N., Caviness, V. S., Jr, … Tsuang, M. T. (2001). Normal sexual dimorphism of the adult human brain assessed by in vivo magnetic resonance imaging. Cerebral Cortex, 11(6), 490–497.

Good, C. D., Johnsrude, I., Ashburner, J., Henson, R. N., Friston, K. J., & Frackowiak, R. S. (2001). Cerebral asymmetry and the effects of sex and handedness on brain structure: a voxel-based morphometric analysis of 465 normal adult human brains. NeuroImage, 14(3), 685–700.

Görgen, K., Hebart, M. N., Allefeld, C., & Haynes, J.-D. (2017). The same analysis approach: Practical protection against the pitfalls of novel neuroimaging analysis methods. NeuroImage. https://doi.org/10.1016/j.neuroimage.2017.12.083

Groen, I. I., Greene, M. R., Baldassano, C., Fei-Fei, L., Beck, D. M., & Baker, C. I. (2018). Distinct contributions of functional and deep neural network features to representational similarity of scenes in human brain and behavior. eLife, 7. https://doi.org/10.7554/eLife.32962

Guggenmos, M., Sterzer, P., & Cichy, R. M. (2018). Multivariate pattern analysis for MEG: A comparison of dissimilarity measures. NeuroImage, 173, 434–447.

Gur, R. C., Turetsky, B. I., Matsui, M., Yan, M., Bilker, W., Hughett, P., & Gur, R. E. (1999). Sex differences in brain gray and white matter in healthy young adults: correlations with cognitive performance. The Journal of Neuroscience: The Official Journal of the Society for Neuroscience, 19(10), 4065–4072.

Haufe, S., Meinecke, F., Görgen, K., Dähne, S., Haynes, J.-D., Blankertz, B., & Bießmann, F. (2014). On the interpretation of weight vectors of linear models in multivariate neuroimaging. NeuroImage, 87, 96–110.

Haxby, J. V. (2012). Multivariate pattern analysis of fMRI: The early beginnings. NeuroImage, 62(2), 852–855.

Haxby, J. V., Gobbini, M. I., Furey, M. L., Ishai, A., Schouten, J. L., & Pietrini, P. (2001). Distributed and overlapping representations of faces and objects in ventral temporal cortex. Science, 293(5539), 2425–2430.

Hebart, M. N., & Baker, C. I. (2017). Deconstructing multivariate decoding for the study of brain function. NeuroImage. https://doi.org/10.1016/j.neuroimage.2017.08.005

Hebart, M. N., Bankson, B. B., Harel, A., Baker, C. I., & Cichy, R. M. (2018). The representational dynamics of task and object processing in humans. eLife, 7. https://doi.org/10.7554/eLife.32816

Jamalabadi, H., Alizadeh, S., Schönauer - Human brain …, M., & 2016. (2016). Classification based hypothesis testing in neuroscience: Below-chance level classification rates and overlooked statistical properties of linear parametric classifiers. Wiley Online Library. Retrieved from http://onlinelibrary.wiley.com/doi/10.1002/hbm.23140/full

Jimura, K., & Poldrack, R. A. (2012). Analyses of regional-average activation and multivoxel pattern information tell complementary stories. Neuropsychologia, 50(4), 544–552.

Joel, D., & Fausto-Sterling, A. (2016). Beyond sex differences: new approaches for thinking about variation in brain structure and function. Philosophical Transactions of the Royal Society of London. Series B, Biological Sciences, 371(1688), 20150451.

Johnstone, T., Ores Walsh, K. S., Greischar, L. L., Alexander, A. L., Fox, A. S., Davidson, R. J., & Oakes, T. R. (2006). Motion correction and the use of motion covariates in multiple-subject fMRI analysis. Human Brain Mapping, 27(10), 779–788.

Kostro, D., Abdulkadir, A., Durr, A., Roos, R., Leavitt, B. R., Johnson, H., … Track-HD Investigators. (2014). Correction of inter-scanner and within-subject variance in structural MRI based automated diagnosing. NeuroImage, 98, 405–415.

Kriegeskorte, N., Mur, M., & Bandettini, P. (2008). Representational similarity analysis - connecting the branches of systems neuroscience. Frontiers in Systems Neuroscience, 2, 4.

Kriegeskorte, N., Simmons, W. K., Bellgowan, P. S. F., & Baker, C. I. (2009). Circular analysis in systems neuroscience: the dangers of double dipping. Nature Neuroscience, 12(5), 535–540.

LaRocque, J. J., Lewis-Peacock, J. A., Drysdale, A. T., Oberauer, K., & Postle, B. R. (2013). Decoding attended information in short-term memory: an EEG study. Journal of Cognitive Neuroscience, 25(1), 127–142.

Long, B., Yu, C. P., & Konkle, T. (2017). A mid-level organization of the ventral stream. bioRxiv. Retrieved from https://www.biorxiv.org/content/early/2017/11/10/213934.abstract

Lüders, E., Steinmetz, H., & Jäncke, L. (2002). Brain size and grey matter volume in the healthy human brain. Neuroreport, 13(17), 2371–2374.

McGrath, J., Saha, S., Chant, D., & Welham, J. (2008). Schizophrenia: a concise overview of incidence, prevalence, and mortality. Epidemiologic Reviews, 30, 67–76.

Naselaris, T., & Kay, K. N. (2015). Resolving Ambiguities of MVPA Using Explicit Models of Representation. Trends in Cognitive Sciences, 19(10), 551–554.

Norman, K. A., Polyn, S. M., Detre, G. J., & Haxby, J. V. (2006). Beyond mind-reading: multi-voxel pattern analysis of fMRI data. Trends in Cognitive Sciences, 10(9), 424–430.

O’Brien, L. M., Ziegler, D. A., Deutsch, C. K., Frazier, J. A., Herbert, M. R., & Locascio, J. J. (2011). Statistical adjustments for brain size in volumetric neuroimaging studies: some practical implications in methods. Psychiatry Research, 193(2), 113–122.

Ojala, M., & Garriga, G. C. (2010). Permutation Tests for Studying Classifier Performance. Journal of Machine Learning Research: JMLR, 11(Jun), 1833–1863.

Parra, L. C., Spence, C. D., Gerson, A. D., & Sajda, P. (2005). Recipes for the linear analysis of EEG. NeuroImage, 28(2), 326–341.

Pedregosa, F., Varoquaux, G., Gramfort, A., Michel, V., Thirion, B., Grisel, O., … Duchesnay, É. (2011). Scikit-learn: Machine Learning in Python. Journal of Machine Learning Research: JMLR, 12(Oct), 2825–2830.

Popov, V., Ostarek, M., & Tenison, C. (2018). Practices and pitfalls in inferring neural representations. NeuroImage, 174, 340-351.

Quionero-Candela, J., Sugiyama, M., Schwaighofer, A., & Lawrence, N. D. (2009). Dataset Shift in Machine Learning. The MIT Press.

Ramakrishnan, K., Scholte, H. S., Groen, I. I. A., Smeulders, A. W. M., & Ghebreab, S. (2014). Visual dictionaries as intermediate features in the human brain. Frontiers in Computational Neuroscience, 8, 168.

Rao, A., Monteiro, J. M., Mourao-Miranda, J., & Alzheimer’s Disease Initiative. (2017). Predictive modelling using neuroimaging data in the presence of confounds. NeuroImage, 150, 23–49.

Ritchie, J. B., Kaplan, D. M., & Klein, C. (2017). Decoding the Brain: Neural Representation and the Limits of Multivariate Pattern Analysis in Cognitive Neuroscience. The British Journal for the Philosophy of Science. https://doi.org/10.1093/bjps/axx023

Rosenblatt, J. D. (2016). Multivariate revisit to “sex beyond the genitalia.” Proceedings of the National Academy of Sciences of the United States of America, 113(14), E1966–E1967.

Sedgwick, P. (2013). Analysing case-control studies: adjusting for confounding. BMJ: British Medical Journal, 346. Retrieved from http://search.proquest.com/openview/29e76b7a6e7e73219e9173cb9eb462bc/1?pq-origsite=gscholar&cbl=2040978

Sepehrband, F., Lynch, K. M., Cabeen, R. P., Gonzalez-Zacarias, C., Zhao, L., D’Arcy, M., … Clark, K. A. (2018). Neuroanatomical morphometric characterization of sex differences in youth using statistical learning. NeuroImage, 172, 217–227.

Smith, S. M., Jenkinson, M., Johansen-Berg, H., Rueckert, D., Nichols, T. E., Mackay, C. E., … Behrens, T. E. J. (2006). Tract-based spatial statistics: voxelwise analysis of multi-subject diffusion data. NeuroImage, 31(4), 1487–1505.

Smith, S. M., Jenkinson, M., Woolrich, M. W., Beckmann, C. F., Behrens, T. E. J., Johansen-Berg, H., … Matthews, P. M. (2004). Advances in functional and structural MR image analysis and implementation as FSL. NeuroImage, 23 Suppl 1, S208–S219.

Smith, S. M., & Nichols, T. E. (2018). Statistical Challenges in “Big Data” Human Neuroimaging. Neuron, 97(2), 263–268.

Snoek, L. (2017). skbold: Utilities and tools for machine learning on BOLD-fMRI data. DOI: 10.5281/zenodo.852473. Available from https://github.com/lukassnoek/skbold.

Todd, M. T., Nystrom, L. E., & Cohen, J. D. (2013). Confounds in multivariate pattern analysis: Theory and rule representation case study. NeuroImage, 77, 157–165.

Van Haren, N. E., Cahn, W., Hulshoff Pol, H. E., & Kahn, R. S. (2013). Confounders of excessive brain volume loss in schizophrenia. Neuroscience and Biobehavioral Reviews, 37(10 Pt 1), 2418–2423.

van Waarde, J. A., Scholte, H. S., van Oudheusden, L. J. B., Verwey, B., Denys, D., & van Wingen, G. A. (2014). A functional MRI marker may predict the outcome of electroconvulsive therapy in severe and treatment-resistant depression. Molecular Psychiatry, 20, 609.

Wacholder, S., Silverman, D. T., McLaughlin, J. K., & Mandel, J. S. (1992). Selection of controls in case-control studies. III. Design options. American Journal of Epidemiology, 135(9), 1042–1050.

Walther, A., Nili, H., Ejaz, N., Alink, A., Kriegeskorte, N., & Diedrichsen, J. (2016). Reliability of dissimilarity measures for multi-voxel pattern analysis. NeuroImage, 137, 188–200.

Weichwald, S., Meyer, T., Özdenizci, O., Schölkopf, B., Ball, T., & Grosse-Wentrup, M. (2015). Causal interpretation rules for encoding and decoding models in neuroimaging. NeuroImage, 110, 48–59.

Woolgar, A., Golland, P., & Bode, S. (2014). Coping with confounds in multivoxel pattern analysis: what should we do about reaction time differences? A comment on Todd, Nystrom & Cohen 2013. NeuroImage, 98, 506–512.

Yu-Feng, Z., Yong, H., Chao-Zhe, Z., Qing-Jiu, C., Man-Qiu, S., Meng, L., … Yu-Feng, W. (2007). Altered baseline brain activity in children with ADHD revealed by resting-state functional MRI. Brain and Development, 29(2), 83–91.

Zhang, Y., Brady, M., & Smith, S. (2001). Segmentation of brain MR images through a hidden Markov random field model and the expectation-maximization algorithm. IEEE Transactions on Medical Imaging, 20(1), 45–57.

